# The C-terminus of the cargo receptor Erv14p affects COPII vesicle formation and cargo delivery

**DOI:** 10.1101/2022.08.13.503845

**Authors:** Daniel Lagunas-Gomez, Carolina Yañez-Dominguez, Guadalupe Zavala-Padilla, Charles Barlowe, Omar Pantoja

**Author notes:** Corresponding author: Omar Pantoja, Instituto de Biotecnología, Universidad Nacional Autónoma de México, Av. Universidad 2001, Cuernavaca, Morelos, 62210, México.

## Abstract

The endoplasmic reticulum (ER) is the start site of the secretory pathway, where newly synthesized secreted and membrane proteins are packaged into COPII vesicles through direct interaction with the COPII coat or aided by specific cargo receptors. Little is known about how post-translational modification events regulate packaging of cargo into COPII vesicles. Erv14/Cornichon belongs to a conserved family of cargo receptors required for the selection and ER export of transmembrane proteins. In this work, we show the importance of a phosphorylation consensus site (Serine-134) at the C-terminus of Erv14. Mimicking phosphorylation of S134 (S134D) prevents the incorporation of Erv14 into COPII vesicles, delays cell growth, exacerbates growth of *sec* mutants, modifies ER structure, and affects localization of several plasma membrane transporters. In contrast, the dephosphorylated mimic (S134A) had less deleterious effects, but still modifies ER structure and slows cell growth. Our results suggest that a possible cycle of phosphorylation and dephosphorylation is important for the correct functioning of Erv14p.

**Summary statement:** Erv14 C-terminus regulates COPII formation and cargo delivery

## Introduction

The early secretory pathway in eukaryotic cells is a process important for the delivery of membrane and soluble proteins to various cellular destinations. Once proteins have been correctly folded and modified in the endoplasmic reticulum (ER), they are sorted from ER-resident proteins and selectively incorporated into COPII-coated vesicles that bud from the ER membrane at ER-exit sites (ERES) to be transported to the Golgi Apparatus (GA) (Brandizzi and Barlowe, 2013), although some proteins exit the ER non-selectively via bulk flow (Martínez-Menárguez et al., 1999; Thor et al., 2009). COPII-coated vesicle generation begins when the GTPase Sar1p is activated by the guanine nucleotide exchange factor (GEF) Sec12, causing its incorporation into ER membranes where it recruits the inner coat layer, Sec24-Sec23 complex. Coat polymerization and vesicle budding occur when Sec23-Sec24 recruits a second complex, the tetramer composed of Sec13-Sec31 that forms the outer coat layer. These events lead to the hydrolysis of GTP on Sar1p and vesicle budding (Sato and Nakano, 2007; Zanetti et al., 2012). The loading of cargo proteins into these vesicles can occur either directly by binding specific COPII subunits, such as Sec24, that binds sorting signals in cargo proteins, or through the participation of proteins called Cargo Receptors (CR), which link cargo proteins indirectly to the COPII-coat (Dancourt and Barlowe, 2010; Herzig et al., 2012; Miller et al., 2002). Among the cargo receptors that have been characterized are members of the p24 family, (Emp24p, Erv25p, Erp1p, and Erp2p) that form the p24 complex and selectively recruits Glycosylphosphatidylinositol (GPI)-anchored proteins (Muñiz et al., 2000) as well as Erv29 or mammalian ERGIC-53 related family members, that transport soluble secreted glycoproteins (Appenzeller et al., 1999; Belden and Barlowe, 2001). The Erv14p/cornichon family are transmembrane proteins that cycle between the ER and early GA, and are involved in the selection of membrane proteins into COPII vesicles (Pagant et al., 2015), most of which are proteins that reside in the plasma membrane (Herzig et al., 2012). Previous reports on the functioning of ScErv14 identified a cytosolic motif (IFRTL) that is required for COPII binding and ER exit of the receptor (Powers and Barlowe, 2002). Subsequent studies have proposed that cargo selection occurs through recognition of long transmembrane domains (TMDs), which are characteristic of plasma membrane proteins (Herzig et al., 2012). It has also been proposed that some cargos bind to residues in the second TMD of Erv14, which together with a new site identified in Sec24p, is proposed to form a dual interaction with both cargo proteins and Erv14. In this view, Erv14p could act as a classical cargo receptor, binding simultaneously to cargo proteins and the COPII coat to drive trafficking (Pagant et al., 2015).

However, little is known about the regulation of the anterograde transport between the ER and the Golgi complex by posttranslational modifications, such as protein phosphorylation (Davis et al., 2016; Koreishi et al., 2013; Lord et al., 2011). In *S. cerevisiae*, it has been reported that some COPII coat subunits are phosphorylated by the serine/threonine kinase Hrr25 (a CK1δ orthologue), allowing coat release and vesicle fusion with the Golgi (Lord et al., 2011). On the other hand, it has been reported that Sit4, a serine/threonine phosphatase, is important for COPII coat dephosphorylation. In the *sit4Δ* mutant, the COPII proteins are hyperphosphorylated and their subcellular locations are modified (Bhandari et al., 2013).

In a previous report we demonstrated the presence and importance of a conserved acidic motif at the C-terminus of plant and yeast Erv14/cornichon homologues, that it is necessary for the binding of the cargo receptor to cargo proteins and their trafficking to the plasma membrane (Rosas-Santiago et al., 2017). Additional analysis of the C-terminus of Erv14 suggested the presence of a motif that could function as a consensus sequence (ESXDD) for casein kinase 2 (CK2). In this work, we show that possible phosphorylation/dephosphorylation of Ser-134 is necessary for the correct packaging of Erv14 into the COPII vesicles and as a consequence, the proper trafficking of its cargo proteins to the plasma membrane.

## Results

### 1.1. A phospho-mimetic state of Ser-134 affects the trafficking of PM proteins

We recently identified an acidic motif (**E**SX**DD**) at the C-terminus of Erv14 that functions as an interaction site with its cargo proteins. This acidic motif is only present in plants and fungi homologues, but not in higher organisms (Rosas-Santiago et al. 2017). To characterize more deeply the role of the C-terminus of Erv14p, we identified a putative phosphorylation site at Ser-134 (Fig. S1). Serine-134 is located within the acidic motif which could function as a consensus sequence (**E**SX**DD**) for casein kinase 2 (CK2) (St-Denis et al., 2015). This observation opened the possibility that the posttranslational modification of Erv14 may play a role in the trafficking of membrane proteins. In order to identify if Serine-134 was phosphorylated, we analyzed the band corresponding to the HA-tagged protein by mass spectrometry (LC–MS/MS), but were unable to identify any protein modification, but confirmed the presence of Erv14 according to the identification of the peptides (IYNKVQLLDATEIFR) and (VQLLDATEIFR) (Fig. S2). In view of these results, we took an alternative approach by generating mutations in Ser-134. We mimicked the phosphorylated and non-phosphorylated states of S134 with the mutations S134D (Erv14^*S134D*^) and S134A (Erv14^*S134A*^), respectively. When we transformed Erv14^*S134A*^ or Erv14^*S134D*^ into the strain BY4741*Δerv14* and monitored cell growth rate, we observed a slower growth compared to the wild-type, and even slower for cells transformed with the empty vector pDR-F1 (Fig. S3). These data indicated the importance of Ser-134 for the correct functioning of Erv14. In view of this evidence and previous reports where Erv14 has been identified as an important cargo receptor, mainly for plasma membrane (PM) proteins (Herzig et al., 2012; Pagant et al., 2015), we tested whether the mutation of Ser-134 modified the targeting of the sodium exchanger Nha1p to the PM. Using the salt-sensitive strain BYT45*erv14Δ* (Navarrete et al., 2010; Rosas-Santiago et al., 2016; Rosas-Santiago et al., 2017), co-expression of Nha1-GFP with the wild type Erv14 or the Erv14^*S134A*^ mutant, showed localization of the exchanger to the PM (Fig. 1A); conversely, the antiporter was retained at the ER upon co-expression with the phospho-mimetic mutant Erv14^*S134D*^ or the empty plasmid pDR-F1 (Fig. 1A). These results indicated that the phosphorylation of Ser-134 prevented the correct traffic of Nha1 to the plasma membrane. To corroborate these results, we next analyzed the localization of the antiporter at the ER (P13) or Golgi/PM (P100) fractions by differential centrifugation. As observed in Figure 1B, when *NHA1-GFP* was co-transformed with wild type *ERV14* or *ERV14^S134A^* into the BYT45*erv14Δ* strain, the antiporter was detected mostly at the PM (P100) fraction, indicative of normal trafficking. In contrast, when Nha1-GFP was co-expressed with the mutant Erv14^*S134D*^, the antiporter remained at the ER (P13) fraction (Fig. 1B). These results were confirmed by quantifying the intensity of the bands which showed a low abundance of Nha1 at the PM fraction relative to the ER in cells co-expressing Nha1-GFP and Erv14^*S134D*^ (Fig. S4). Thus, demonstrating that the phosphorylated mimetic of Ser-134 prevents the correct trafficking of its client. Additional confirmation on the importance of the phosphorylation state of Erv14 for the correct targeting of Nha1 to the PM was derived from assessing the sensitivity of the yeast cells to sodium. Co-expression of Nha1-GFP and Erv14 or the Erv14^*S134A*^ mutant, rescued the tolerance to sodium to *BYT45erv14Δ* cells, in contrast to cells co-transformed with *NHA-GFP* and *ERV14^S134D^* which were more sensitive to the cation (Fig. 1C). Together, these results are indicative of an impaired delivery of Nha1 to the plasma membrane by the Erv14^*S134D*^ mutant.

**Figure 1.**
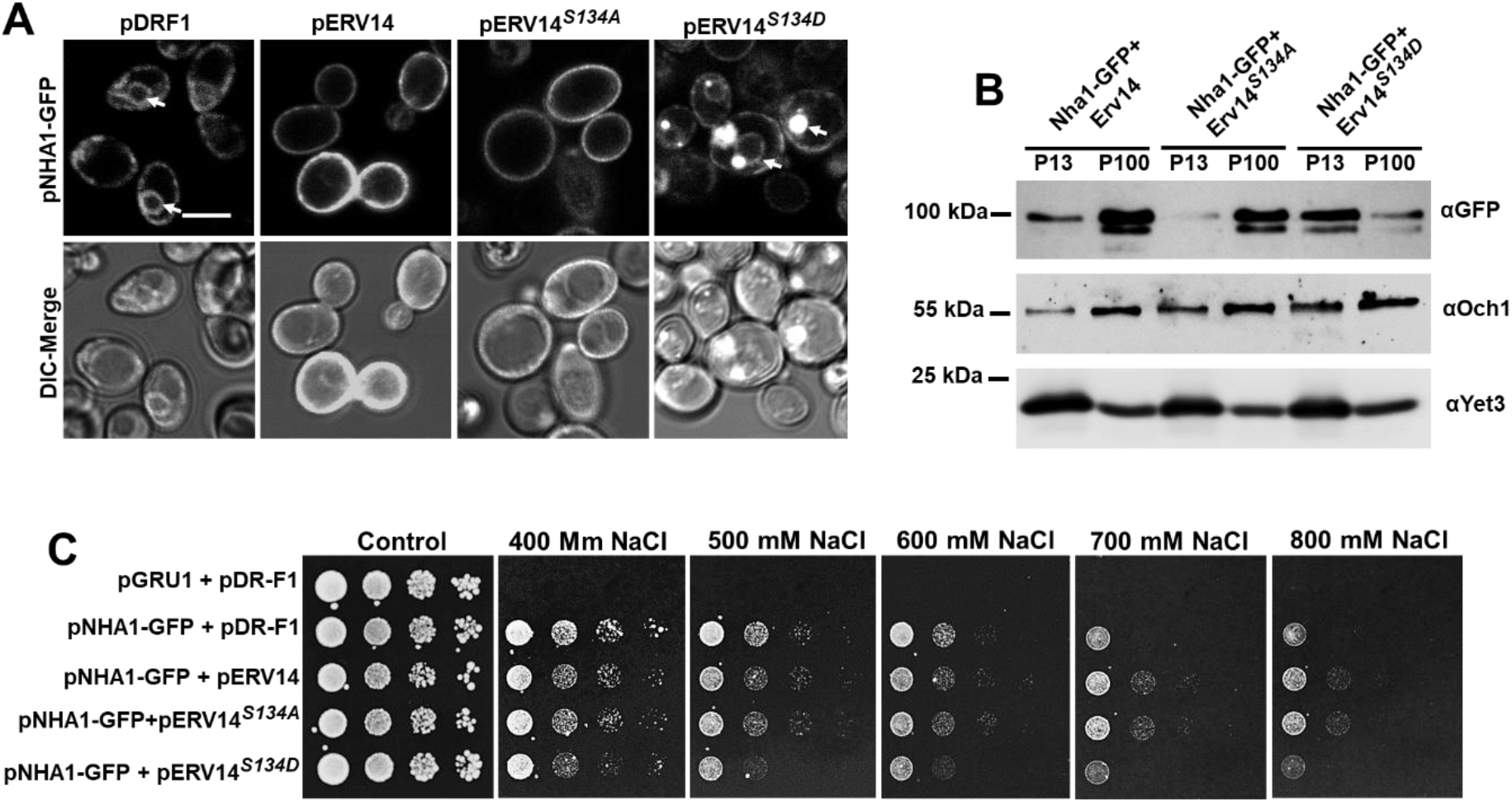
The phosphomimic mutation S134D in Erv14 affects the targeting of the Nha1 exchanger to the plasma membrane. (**A**) Plasma membrane localization of the antiporter Nha1-GFP was observed in BYT45*erv14Δ* cells transformed with wild type pERV14 or pERV14^*S134A*^, but not with pDR-F1 (empty) or *pERV14^S134D^*. Arrows indicate perinuclear ER. (Scale bar, 2 μm). (**B**) Confirmation of proper localization of Nha1 to the PM by pERV14 or pERV14^*S134A*^, but not by pERV14^*S134D*^. Och1 and Yet3 were used as markers for the Golgi apparatus and endoplasmic reticulum, respectively. (**C**) Salt sensitivity of the BYT45*erv14Δ* cells was rescued by co-transformation with pNHA1-GFP and pERV14 or pERV14^*S134A*^, but not by co-transformation with pERV14^*S134D*^.

Erv14 has been identified as a cargo receptor that is required for efficient ER export of several proteins, a vast majority of which locate to the plasma membrane (Herzig et al., 2012; Pagant et al., 2015; Rosas-Santiago et al., 2017). To analyze if more clients whose transport out of the ER is dependent on Erv14 were affected by the phosphorylation state of the receptor, we examined Pdr12, a weak acid-efflux ATPase from the ABC family (Piper et al., 1998) and Qdr2, a drug/proton antiporter of the MFS family (Vargas et al., 2004). The co-transformation of BY4741*pdr12Δerv14Δ* cells with *PDR12-GFP* and *ERV14* or *ERV14^S134A^*, did not affect the delivery of the Pdr12-GFP to the plasma membrane (Fig. 2A). In comparison, co-expression of Pdr12-GFP with Erv14^*S134D*^ or the empty pDR-F1 vector caused retention of the ABC transporter at the ER, as clearly shown by the fluorescence signal around the nucleus and the cell periphery, corresponding to the perinuclear and cortical localization of the ER in yeast cells (Fig. 2A). Similar results were obtained with Qdr2, where we observed correct targeting of the exchanger to the PM with either Erv14 or Erv14^*S134A*^ (Fig. 2B) but not with Erv14^*S134D*^ or the empty vector (Fig. 2B). These results confirmed that the phosphomimetic state of Erv14 controls the traffic of its cargo proteins.

**Figure 2.**
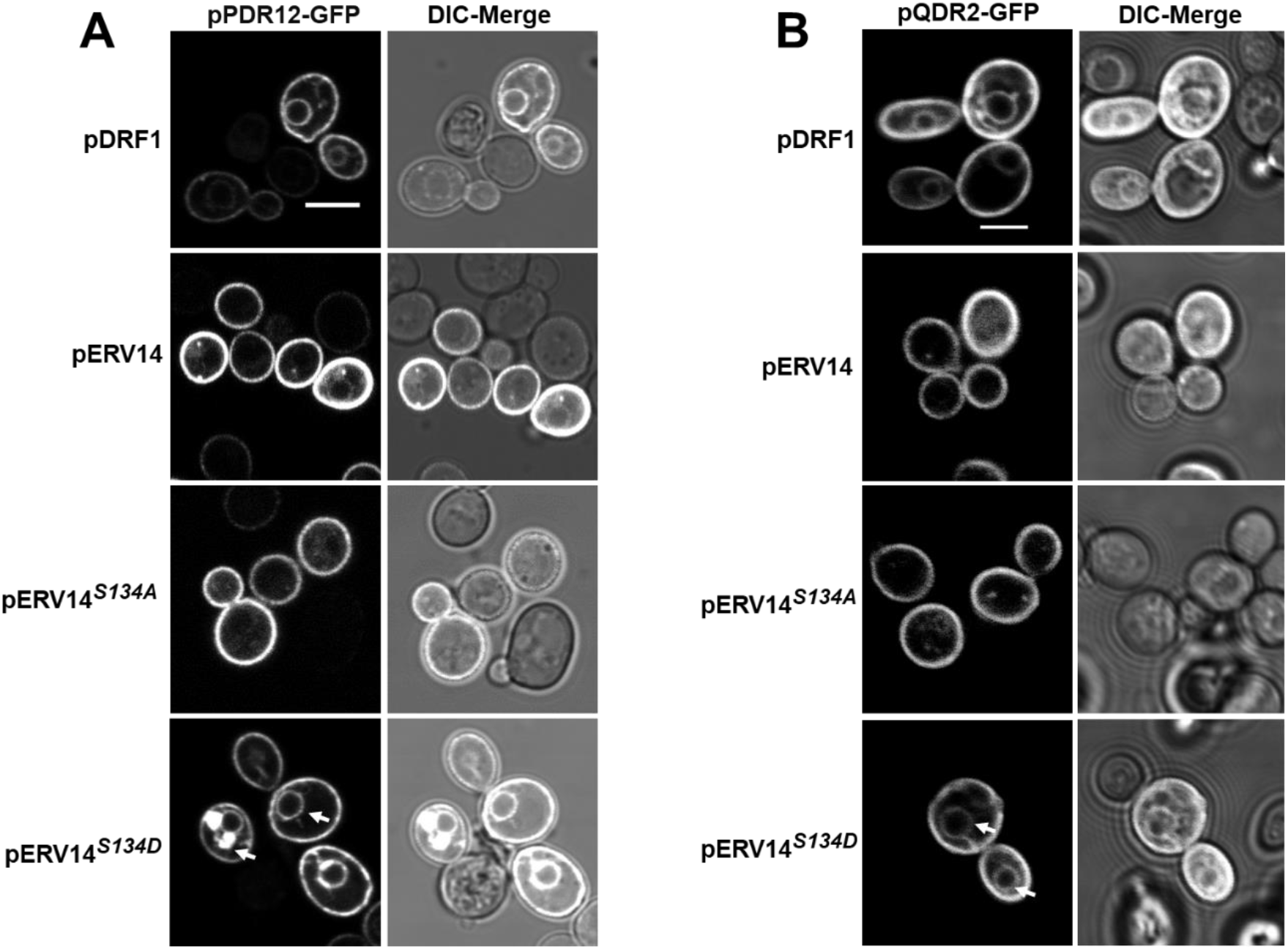
The S134D mutation at the C-terminus in Erv14 prevents export of plasma membrane cargos. Trafficking to the plasma membrane of the ABC transporter Pdr12-GFP or the MFS protein Qdr2-GFP in the BY4741*pdr12Δerv14Δ* (**A**) or BY4741*qdr2Δerv14Δ* (**B**) cells only occurred in cells co-transformed with wild type pERV14 or pERV14^*S134A*^, but not with the empty vector (pDR-F1) or pERV14^*S134D*^, where cargo proteins showed a perinuclear ER retention (arrows). (Scale bar, 2 μm).

### 1.2. Ser-134 is not required for interaction with the cargo proteins

So far, our results demonstrate that the phospho-mimic form impaired delivery of cargo proteins to the plasma membrane. We next reasoned that the potential role for the phosphorylation state of Ser-134 in Erv14 could control or function as a cargo binding site. We sought to test this hypothesis by monitoring Erv14-cargo interaction using the membrane-based Split-Ubiquitin Yeast Two Hybrid System (mbSUS) (Stagljar et al., 1998). For this, we employed Qdr2 as bait (Cub clone), which we had previously identified as interacting with Erv14 (Rosas-Santiago et al., 2017), and the point mutants at Ser-134 as prey (Nub clone). Taking advantage of the repressible Met promoter present in the Cub constructs, we titrated the strength of the interaction employing increasing concentrations of Met (Grefen et al., 2009). Erv14 and Qdr2 showed robust interaction, as indicated by the growth of diploid cells in the selection medium (Ura-, Trp-, Leu, Ade-, His-) (Fig. 3A). The growth of diploid cells was not affected by any of the two mutations, either Erv14^*S134A*^ or Erv14^*S134D*^, indicating that modifications at Ser-134 did not influence the interaction between receptor and cargo. Moreover, the strength of these interactions was similar, as indicated by the similar inhibition of cell growth caused by the increasing concentrations of Met (Fig. 3A). To confirm the interaction of Qdr2 and the Erv14 isoforms, we tested the activation of the *lacZ* gene by incubating the cells in presence of X-Gal. As observed in Figure 3B, the development of a blue signal in diploid cells expressing Qdr2 and wild-type Erv14 or mutants forms, was of similar intensity, suggesting a similar interaction between the three Erv14 isoforms and the Qdr2 cargo. Together, these results suggest that Ser-134 does not play an important role in the interaction between the Erv14 cargo receptor and its cargos.

**Figure 3.**
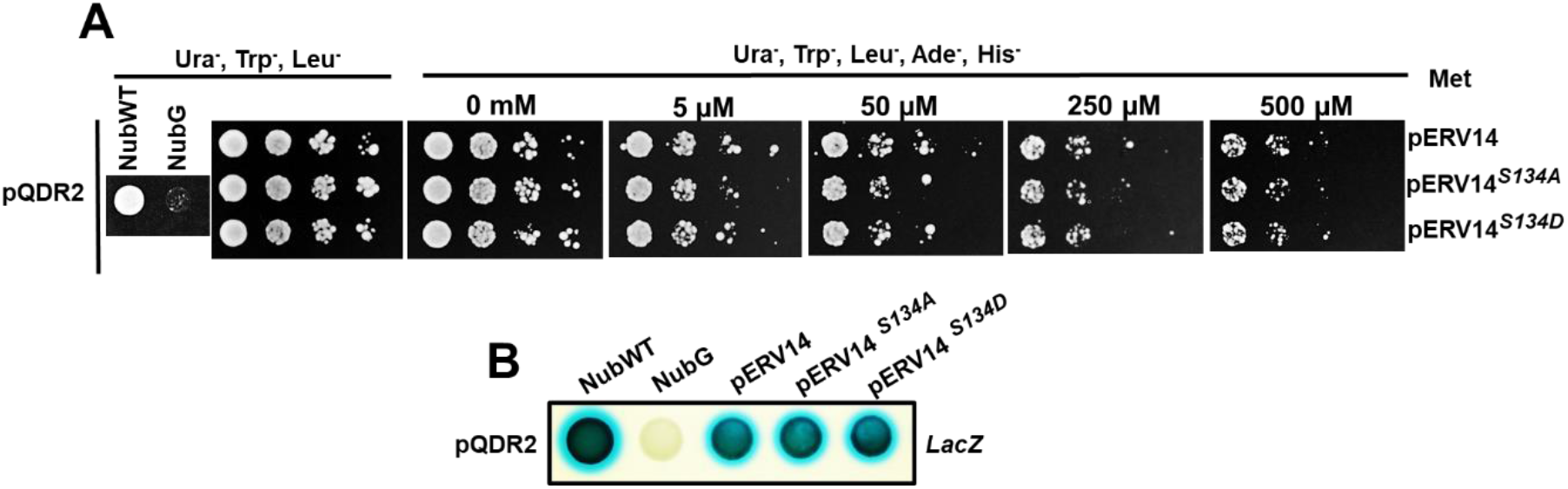
Mutations in Ser-134 do not affect binding with cargo proteins. (**A**) Interaction between Qdr2 and Erv14 isoforms was not affected by the mutation of Ser-134 and showed similar strength at increasing concentrations of methionine. (**B**) Corroboration of the interactions was demonstrated by activation of LacZ and revealed with X-Gal as substrate (LacZ). NubWT and NubG were used as false negative and false positive controls, respectively. Data are representative of at least three repetitions.

### 1.3. Mutation of Ser-134 modified the subcellular location of Erv14 and induced structural changes in the ER

Erv14 is a transmembrane protein that localizes to the ER and Golgi compartments, and is proposed to cycle between these two organelles through the COPII and COPI systems (Powers and Barlowe, 1998; Powers and Barlowe, 2002). In view of our previous results, we wondered if mistargeting of Erv14p cargos could be explained by alterations in localization of the receptor. To analyze this possibility, we generated GFP fusions at the C-terminus of Erv14 and the Ser-134 mutants to follow their localization in vivo by confocal microscopy. Erv14-GFP was found mainly in both, the cortical and the perinuclear ER, with some dots that could be associated to the Golgi apparatus, as previously observed (Rosas-Santiago et al., 2017) (Fig. 4A). The subcellular localization of the mutants Erv14^*S134A*^ or Erv14^*S134D*^ seemed to be like that observed for the wild-type receptor (Fig. 4A). When we monitored trafficking of Erv14p between the ER and Golgi membranes of the wild-type Erv14 and Ser-134 mutants by membrane fractionation and Western blot, the abundance of wild-type Erv14p was similar in the ER and the Golgi fractions, indicative of normal trafficking (Fig. 4B). Conversely, localization of the phosphomimetic Erv14^*S134D*^ mutant was shifted to the ER fraction (P13), indicating its retention at the ER (Fig. 4B). For the Erv14^*S134A*^ mutant, no clear change in membrane distribution was observed (Fig. 4B). The unchanged distribution of the Golgi resident Och1 and the ER marker Yet3 (Fig. 4B), confirmed that the changes observed for the Erv14^*S134D*^ mutant were specific. Additional confirmation for these results was obtained by quantifying the band intensities from the Western blots. For this, the ratios for each protein in the ER (P13 fraction) with respect to the total protein (ER+GA; P13+P100) were quantified, showing that the ratio for the mutant Erv14^*S134D*^ was 20% higher than that for Erv14 (Fig. 4C). For the S134A mutant, a smaller ratio was observed, but it was not significant (Fig. 4C). Based on these results, we explored possible changes caused by the mutations of Erv14 at the ultrastructural level. The wild-type (Erv14) cells showed a clear continuous cortical ER at the TEM level (Fig. 5A). In comparison, cells expressing Erv14^*S134A*^ or Erv14^*S134D*^ showed a deformed cortical ER, with omega-like structures distributed along this organelle (Fig. 5C-D). These defects were not observed in the mutant strain *erv14Δ*, which showed a separation of the cortical ER from the cell’s periphery (Fig. 5B). These structural changes at the ER membrane caused by mutation of Ser-134 could be due to modifications in trafficking between ER and Golgi apparatus, as has been observed for early *sec* mutants that impair vesicle formation (Kaiser and Schekman, 1990; Miller et al., 2002).

**Figure 4.**
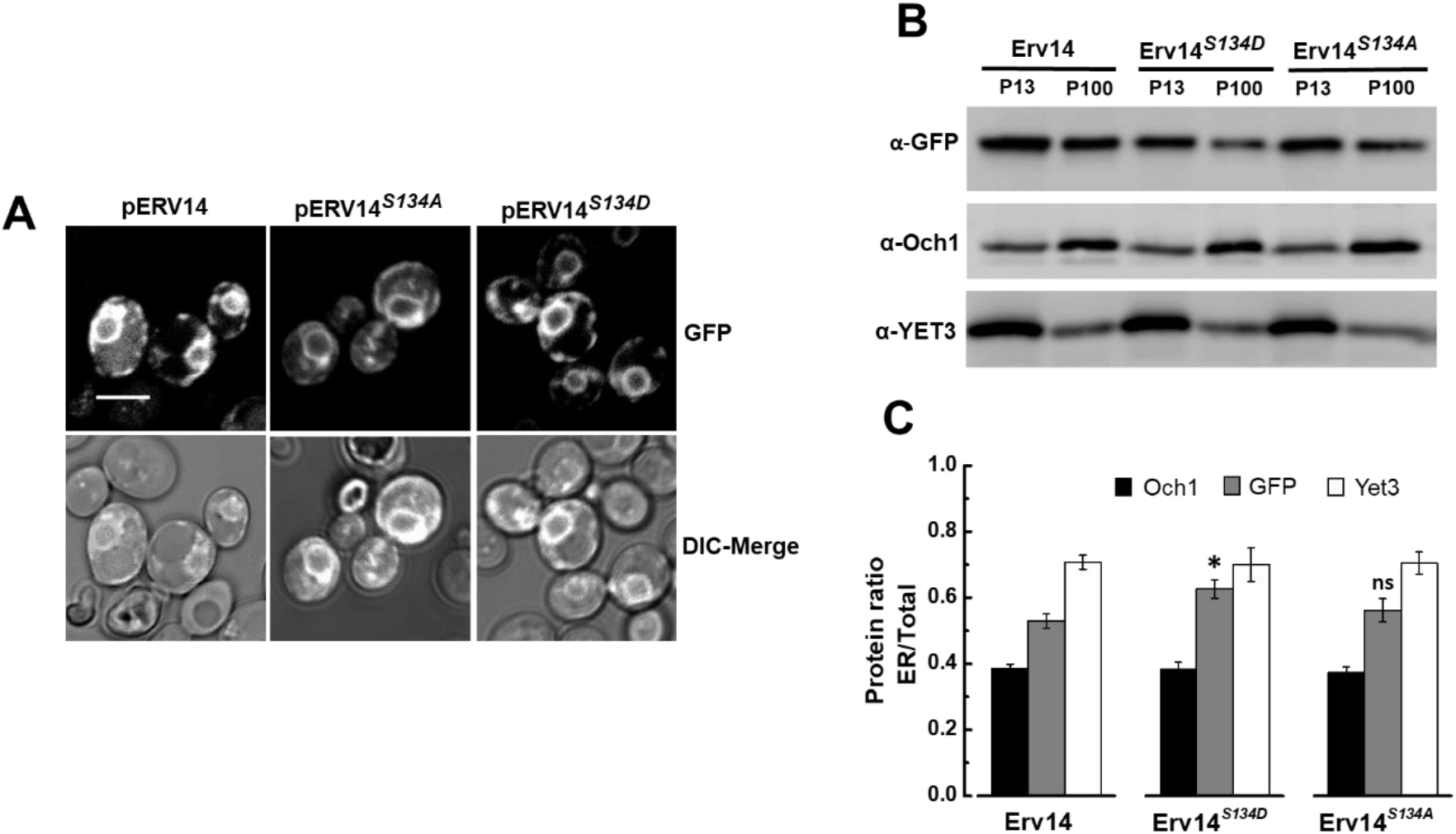
The S134D mutation modified the subcellular location of Erv14. (**A**) GFP-tagged Erv14 isoforms were expressed in the BY4741*erv14Δ* strain. ER and AG membrane localization of GFP-tagged Erv14, Erv14^*S134A*^ or Erv14^*S134D*^ showed similar subcellular localization. (Scale bar, 2 μm). (**B**) Erv14 membrane distribution in ER or Golgi fractions from GFP-tagged wild-type and Erv14^*S134A*^ isoforms was similar, observing a decrease of Erv14^*S134D*^ at the Golgi fraction. Yet3 and Och1 were used as markers for the ER and GA, respectively. (**C**) Bar plot shows the ratio of each protein at the ER against the total (ER+AG). Data are the mean ± SD from three different preparations; The t-test Student analysis was used to compare each mutant to wild type with *p* values of: ns, *p*>0.05, * p<0.05, ** p<0.001.

**Figure. 5.**
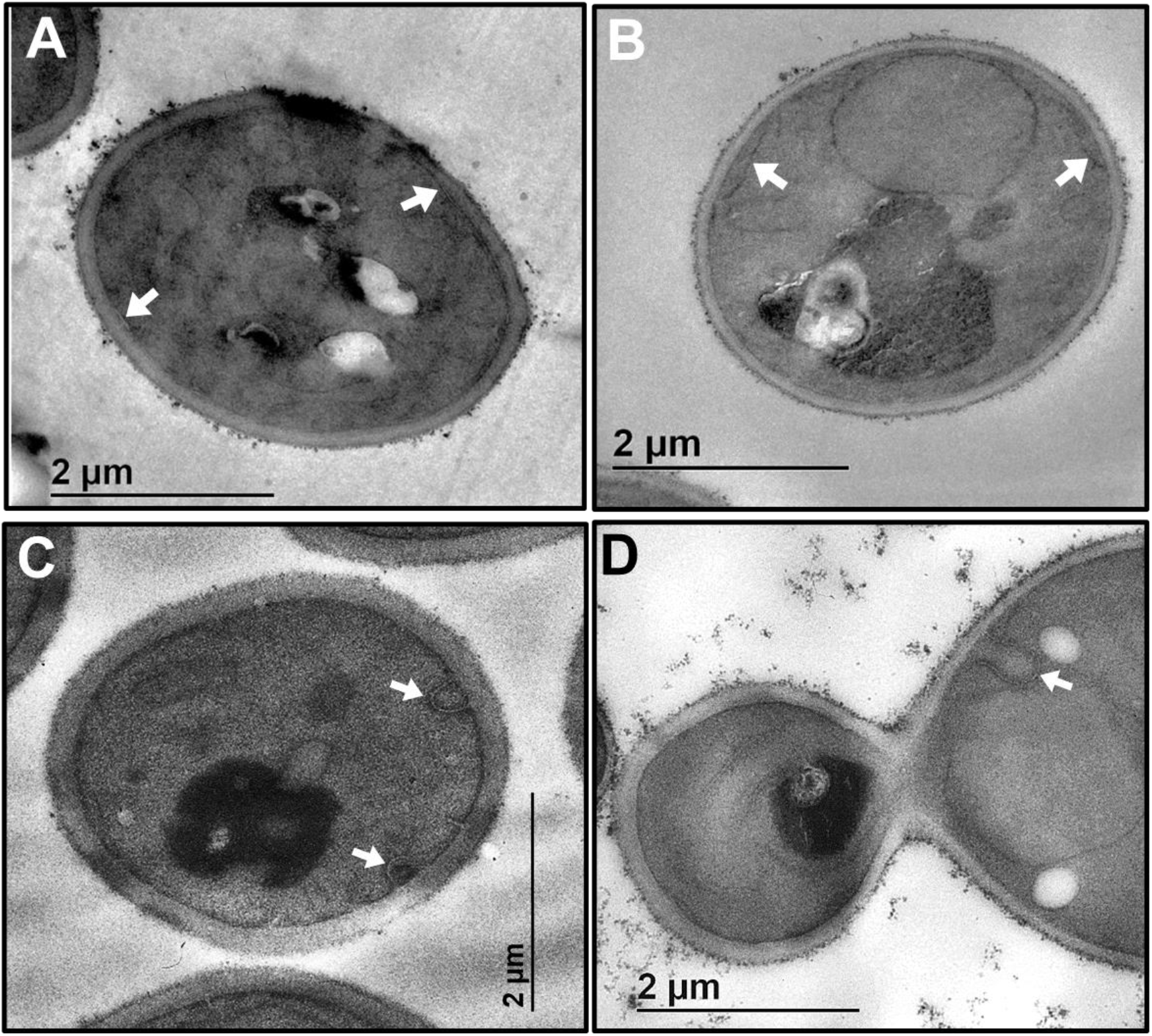
Mutations at Ser-134 induced modifications in ER structure. TEM micrographs from BY4741*erv14Δ* cells expressing Erv14 or the mutants Erv14^S134A^, or Erv14^*S134D*^. (**A**) In Erv14p expressing cells the cortical ER was observed as a continuous thread closely attached to the PM (arrows). (**B**) In mutant *erv14Δ* cells the cortical ER separated from the PM. In cells expressing the mutants Erv14^*S134A*^ (**C)** or Erv14^*S134D*^ (**D**), cortical ER showed deformation in omega like structures (arrows).

### 1.4. The phospho-mimetic form of Erv14 impaired the formation of COPII vesicles

Export of secretory proteins from the ER is mediated by COPII complexes on the ER membrane, driving the formation and budding of COPII coated vesicles (Barlowe et al., 1994). Erv14 was identified as being enriched in COPII vesicles and necessary for the incorporation of cargo proteins into COPII vesicles (Pagant et al., 2015; Powers and Barlowe, 1998; Powers and Barlowe, 2002). Therefore, we asked whether packaging of Erv14 into COPII vesicles was influenced by mutation of the Ser-134 residue using an in vitro assay that recapitulates COPII-dependent vesicle formation (Barlowe et al. 1994). For this, we employed washed semi-intact *BY4742erv14Δ* cells (SICs) expressing HA-tagged Erv14 or the point mutants Erv14^*S134A*^ or Erv14^*S134D*^, incubated with or without purified COPII proteins (Sar1, Sec23/Sec24 and Sec13/Sec31), GTP and an ATP regeneration system to drive vesicle budding (Baker et al., 1998; Barlowe et al., 1994). The vesicle fraction was separated from bulk membranes by centrifugation and ten percent of the total membrane input (T) was compared with the sample of the vesicular fraction by immunoblotting. In SICs expressing Erv14 or Erv14^*S134A*^, Erv14 was efficiently packaged into the COPII vesicles, as indicated by the strong signal developed by the anti-HA antibody (Fig. 6A). However, for vesicles isolated from the SICs expressing Erv14^*S134D*^, packing of Erv14 into COPII vesicles was less efficient, according to the weaker anti-HA signal (Fig. 6A). In all three cases, Erv14 packing did not occur in the absence of COPII proteins (Fig. 6A). As a negative control, we used Sec61, an integral membrane protein that resides in the ER and is a subunit of the translocon, which was not detected in the vesicle fraction (Fig. 6A). In contrast, Coy1, a well characterized membrane protein, was correctly sorted to COPII vesicles (Fig. 6A), that also served as positive control (Anderson et al., 2017; Otte et al., 2001). Unexpectedly, a weaker signal was observed for Erv41 (Fig. 6A). These results were confirmed by quantifying budding efficiency as described in M&M (Fig. 7B). ER-derived vesicles incorporated Erv14 or Erv14^*S134A*^ at 9% and 10%, respectively (Fig. 6B), comparable to results previously reported for Erv14 (Powers and Barlowe, 1998). Conversely, Erv14^*S134D*^ was packed at lower level (4%), (Fig. 6B). On the other hand, when the budding for Erv41 was quantified in membranes expressing Erv14 or Erv14^*S134A*^ we obtained a level of 10% and 11% efficiency, respectively (Fig. 6B). For the membranes expressing Erv14^*S134D*^, packing of Erv41 was less efficient showing a level of 4% (Fig. 6B). These values are similar to those reported for the *ΔErv14* deletion strain, where the transport efficiency of gpαF was also reduced when compared to the wild type (Powers and Barlowe, 2002). Together, these results indicate that the phospho-mimetic mutation at the C-terminus of Erv14 decreases formation of COPII vesicles.

**Figure. 6.**
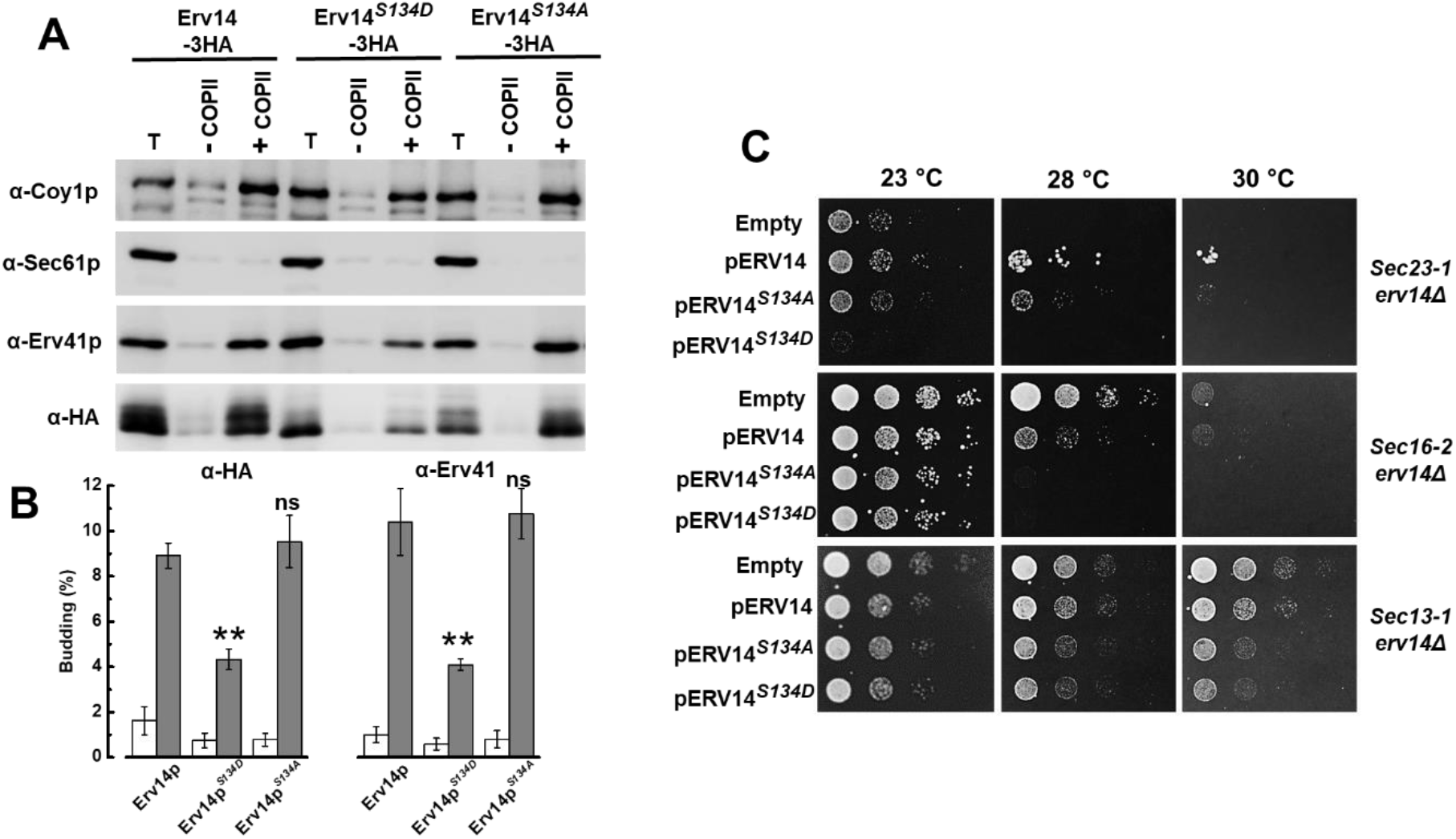
Mutations in Serine-134 affected packaging of Erv14 into COPII vesicles and prevented the rescue of thermosensitive *sec erv14* mutants by *ERV14*. (**A**) Semi-intact BY4742*erv14Δ* cells expressing the wild-type cargo receptor (Erv14-HA), or the mutants (Erv14^*S134A*^-HA) or (Erv14^*S134D*^ HA) were incubated with (+) or without (-) purified COPII proteins to monitor vesicle budding. One tenth of the total budding reaction (T) was included as a control. Budded vesicles were isolated and resolved on a 14% polyacrylamide gel and immunoblotted with antibodies specific for Erv41, Coy1 (positive control), Sec61 (negative control) and HA. (**B**) Quantification of COPII packaging efficiency from three essays as in **A** without (open bars) or in the presence of COPII proteins (solid bars). Vesicle packaging was evaluated with α-HA for Erv14 (left) or α-Erv41 (right). The t-test Student analysis was used to compare each mutant to wild type with *p* values of: ns, *p*>0.05, * p<0.05, ** p<0.001. (**C**). Cell growth in thermo-sensitive *sec23-1, sec16-2* and *sec13-1* cells that was rescued by *pERV14* did not occur with the Erv14^*S134A*^ or Erv14^*S134D*^ mutants. Cells were grown at permissive temperature (23°C), diluted and plated, and grown at either permissive (23°C) or restrictive temperatures (28–30°C). The transformants were grown for 3 d. Representative results from three independent assays.

**Figure 7.**
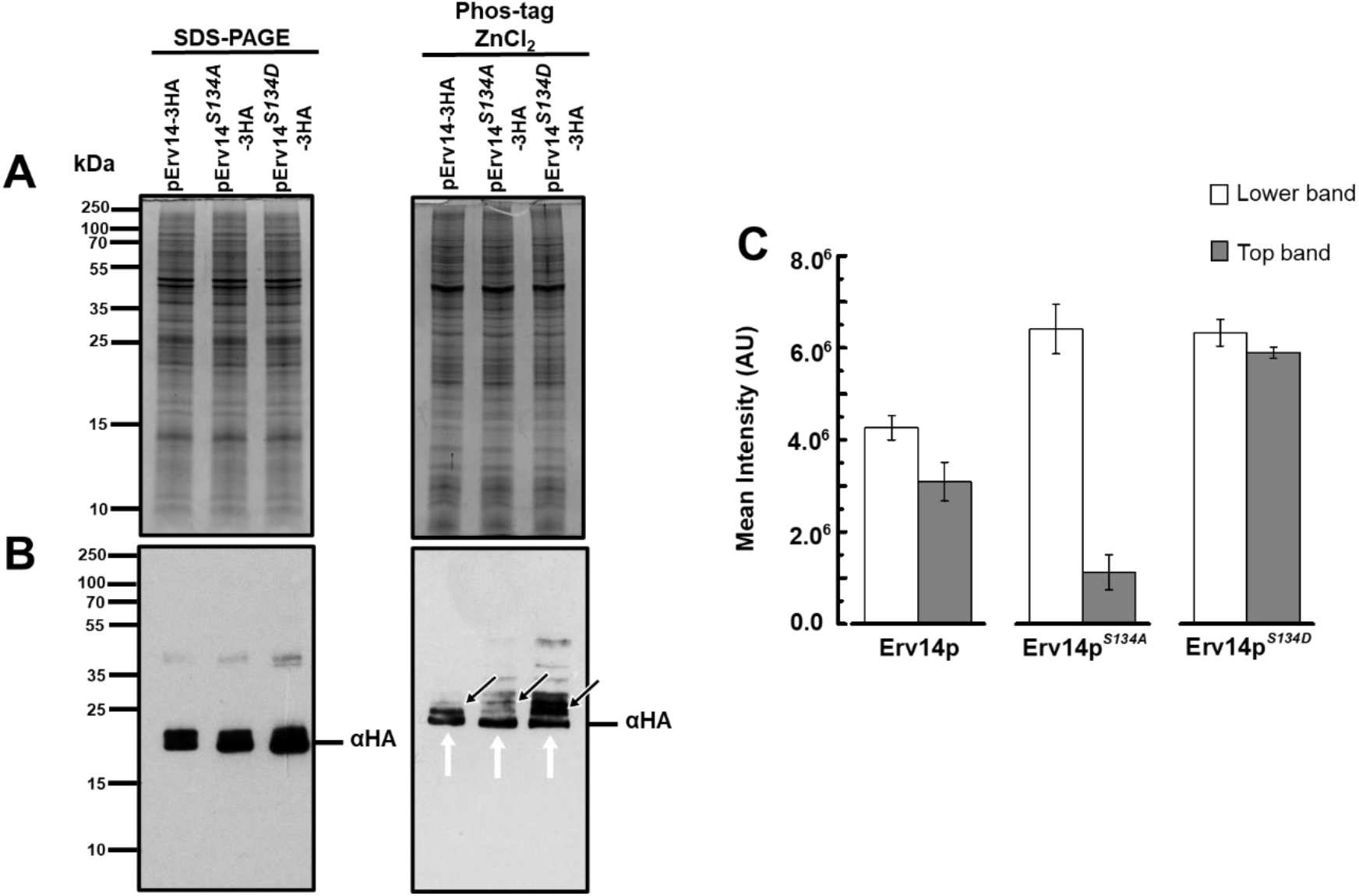
Erv14 is phosphorylated at the carboxyl terminus. (**A**) Microsomal proteins (P13) from HA-tagged Erv14, Erv14^*S134A*^ or Erv14^*S134D*^ expressing cells were separated in SDS-PAGE (left) or Zn(II)-Phos-tag (right) gels. (**B**), HA-tagged Erv14 proteins were identified an α-HA antibody. Arrows indicate phosphorylated (black) and non-phosphorylated (white) isoforms, respectively. (**C**). Bar plot shows the ration intensity from the top to that of the lower band corresponding to the recognition by anti-HA antibody from HA-tagged Erv14 (white bar), Erv14^*S134A*^ (Blackbar) and Erv14^*S134D*^ (dark grey bar) expressing cells. Data are the mean from three different preparations.

### 1.5. Mutation of Ser-134 Erv14 exacerbates the Growth of Secretory Yeast Mutants

In a previous report it was found that deletion of *ERV14*, when combined with temperature sensitive mutations involved in COPII vesicle formation, such as Sec23, Sec16 and Sec31, increased the thermosensitivity of the cells, which indicated that Erv14 was involved in formation of the COPII vesicles (Powers and Barlowe, 2002) (Fig. S5). To support the results of *in vitro* vesicle budding, we examined whether the Erv14 mutants modified the viability of the temperature-sensitive mutants in the early secretory pathway. We placed *ERV14* and the S134 mutants under the control of the constitutive GADP promoter and introduced each construction into the double mutants *sec23-1 erv14Δ, sec13-1 erv14Δ* and *sec16-2 erv14Δ* and grew them at increasing temperatures of 23°C, 28°C and 30°C. As shown in figure 6C, the *sec23-1 erv14Δ* strain transformed with *ERV14^S134D^* did not grow at the semi-restrictive temperature of 28°C nor at 30°C, like the cells transformed with the empty vector. In contrast, *sec23-1 erv14Δ* cells showed a clear growth at 23°C when transformed with *ERV14^S134A^*, although not as well as cells transformed with *ERV14* (Fig 6C). For the *sec16-2 erv14Δ* strain, transformation with *ERV14* reduced cell growth at the non-permissive temperatures of 28°C and 30°C, a response that was exacerbated when the cells were transformed with *ERV14^S134A^* or *ERV14^S134D^* (Fig. 6C). With the *sec13-1 erv14Δ* double mutant, we observed low growth of cells transformed with *ERV14^S134A^* or *ERV14^S134D^;* conversely, when transformed with *ERV14*, cells grew better and similar to those transformed with the empty plasmid (Fig. 6C). Thus, these genetic interactions show that the cells expressing the *ERV14^S134D^* mutant, in particular, inhibited cell growth and corroborated our in vitro budding results that indicate the S134D mutation in Erv14 influences COPII vesicle formation.

### 1.6. Possible phosphorylation of Erv14 at the carboxyl terminus

In view of the previous results, we attempted to identify if Erv14 was phosphorylated using the ZnCl_2_-Phos-tag SDS-PAGE (Wako Chemicals, USA) technique, which is an electrophoresis approach that allows the separation of phosphorylated and non-phosphorylated proteins using conventional SDS-PAGE procedures (Kinoshita and Kinoshita-Kikuta, 2011). For this, we examined possible changes in the migration patterns of the three Erv14 protein versions. As shown in figures 7A, the total protein profile between the cells expressing Erv14, Erv14^*S134A*^ or Erv14^*S134D*^ mutants, was very similar on SDS-PAGE or the Zn^+2^-Phos-tag gel, but different between the two separation matrixes, indicating the modified migration of many proteins in the latter (Fig. 7A). To identify modifications on the mobility of Erv14, Erv14^*S134A*^ or Erv14^*S134D*^ on the two separation matrixes, we analyzed their migration by Western blot, In the conventional SDS-PAGE gel, no difference was observed in the banding pattern between the Erv14 and the mutants Erv14^*S134A*^ or Erv14^*S134D*^ (Fig. 7B). In comparison, we observed in the Zn^+2^-Phos-tag gel the presence of two Erv14 isoforms, with the top one probably corresponding to the phosphorylated form (Fig. 7B). For the Erv14^*S134A*^ and Erv14^*S134D*^ mutants these two bands showed different patterns, with a marked decreased in the signal for the top band from Erv14^*S134A*^, whereas a stronger signal was associated with Erv14^*S134D*^ (Fig. 7B). Additional bands were revealed by the anti-HA antibody, with an apparent increase in the abundance in Erv14^*S134D*^ (Fig. 7B). To confirm these results, the ration between the intensity of the top to that of the lower band was quantified from the Zn^+2^-Phos-tag gel (Fig. 7C). This analysis showed that the amount of the top isoform in Erv14 was 35% less abundant than the lower band a condition that clearly changed for the Erv14^*S134A*^ mutant, where the top band almost disappeared (18%) (Fig. 7C). For the Erv14^*S134D*^ mutant the abundance of both isoforms was very similar (92%) (Fig. 7), suggesting the possible phosphorylation of Erv14 at Ser-134.

## Discussion

Traffic control of membrane proteins from their place of synthesis, the ER, to their place of residence, is tightly regulated by proteins called cargo receptors. The Erv14p family of proteins has been identified as cargo receptors that facilitate the packaging of membrane proteins into COPII-coated vesicles for traffic out of the ER (Herzig et al., 2012; Pagant et al., 2015; Powers and Barlowe, 1998; Powers and Barlowe, 2002). Yet, little is known about the regulation of these cargo receptors by posttranslational modifications. Recently, we showed that the (**E**SG**DD**) acidic motif at the C-terminus of Erv14p is required for interaction and correct targeting of its cargo proteins (Rosas-Santiago et al., 2017). After further analysis, we identified that the C-terminus possesses a consensus sequence for the activity of casein kinase II (CKII) to phosphorylate S134 according to the Netphos3.1 motif prediction program (Fig. S1). This domain agrees with empirical data that demonstrated a required presence of an acidic residue at the position located three residues to the carboxy terminus of the phosphate acceptor site, together with the enhancing effect generated by the presence of an acidic residue on the amino side of the Ser residue, which is preferred by CKII, rather than threonine (Meggio and Pinna, 2003; Salvi et al., 2009; St-Denis et al., 2015). This evidence opened the possibility that the phosphorylation/dephosphorylation of Ser-134 could modify the functioning of Erv14.

By employing mutant analogs for the phosphorylated or dephosphorylated residue Ser-134 we demonstrate that functioning of the Erv14 cargo receptor is affected differentially by both modifications. While Erv14 membrane localization, ER ultrastructure and interaction with the cargos were similarly modified by both mutations, only the phosphomimic S134D mutation significantly reduced the delivery of cargo proteins to the plasma membrane that was associated with subtle changes in the distribution of the receptor between the ER and the GA. These results suggest that the mutation S134D alters the trafficking of its clients through a mechanism that does not seem to involve a direct protein-protein interaction. This observation contrasts with the direct prevention of the interaction of the receptor with its cargos when the acidic amino acids, that form part of the phosphorylation domain, were mutated (Rosas-Santiago et al., 2017).

Our experiments from *in-vitro* assays that reproduced COPII budding, reinforced the view that the mutant Erv14^*S134D*^ affects the packaging of Erv14 into COPII vesicles and also decreases their formation, which could explain the mistargeting of cargo proteins (Fig. 6). The reduced packaging of Erv41 and Coy1, a cargo receptor and a Golgi-located protein, respectively, that are known to travel or use the COPII system (Anderson et al., 2017; Otte et al., 2001), confirms that the phospho-mimetic mutation in Ser-134 affects the formation of the COPII vesicles (Fig. 6). Supporting this observation are the genetic experiments that showed that the expression of Erv14^*S134D*^ exacerbated cell growth of temperature sensitive mutants directly involved in COPII formation (SEC23, SEC16, SEC13) (Fig. 6C). Particularly, the strong inhibition of cell growth of the Sec23 mutant can be used to propose that as Erv14 directly interacts with the Sec23/24p complex (Pagant et al., 2015; Powers and Barlowe, 2002), it may be possible that mimicking phosphorylation of Ser-134 at the C-terminus, inhibits/alters binding of Erv14 to COPII subunits, thus interrupting cargo trafficking.

According to the cornichon molecular model (Nakagawa, 2019) and the one we derived for Erv14 using the Alphafold2 algorithm https://www.deepmind.com (Fig. S6), we can confirm that the C-terminus faces the ER lumen, and the IFRTL domain that has been shown to be important for the interaction with the Sec23/Sec24 complex, faces the cytoplasm (Pagant et al., 2015; Powers and Barlowe, 2002). It is possible that by simulating phosphorylation of S134 a conformational change in Erv14 is induced, precluding its correct interaction with the Sec23/Sec24 complex, and therefore, the formation of COPII vesicles.

According to the changes observed in the TEM images and the genetic interactions with the *sec* mutants, both isoforms, Erv14^*S134D*^ and Erv14^*S134A*^ caused deformations in the ER and worsen cell growth (Fig. 5–6), suggesting an alteration in the ER-to-Golgi transport, as has been observed in several *sec* mutants (Kaiser and Schekman, 1990; Miller et al., 2002).

Indications that Ser134 is phosphorylated were obtained by analysis of the changes on mobility of the modified protein employing the Zn^+2^-Phos-tag technique, where the presence of a second band with a retarded movement was observed for Erv14 (Fig. 7). This observation was confirmed by a decrease in abundance of the slower band in the S134A mutant, while observing an increase in the S134D mutant (Fig. 7). Presence of the top band in the S134D mutant confirm that the mutation seems to be a good representation of the phosphorylated state of Erv14, evidence that help us to conclude that phosphorylation/dephosphorylation of Erv14 is an important mechanism in the functioning of COPII in the transport of plasma membrane proteins. The intensity of the bands associated to Erv14 indicate that the abundance of the phosphorylated protein seems to be slightly lower at the ER, that together with the results from the S134A and S134D mutants, indicate that an alteration in this proportion can have severe consequences in the functioning of the cargo receptor and accordingly, in the formation of COPII vesicles. The detection of additional bands with the S134D mutant may be due to a strong interaction between the carboxylate group from aspartate, and the Zn^+2^-Phos-tag complex. We attempted to confirm these results by analyzing possible changes in the Erv14 isoforms in the absence of CK2, that unfortunately, did not show a clear disappearance of the phosphorylated Erv14 isoform, although a clear decrease in the abundance of the receptor protein was observed in the *Scck2Δ* mutant (Fig. S7). These results suggest that an alternative kinase may be responsible for the phosphorylation of Erv14.

Our results raise the question of where the phosphorylation of Erv14 occurs. It has been reported that numerous proteins are phosphorylated by unidentified protein kinases present within the lumen of the Golgi, enzymes that are poorly characterized and have been termed secreted protein kinases (Park et al., 2019). Moreover, both Sec23 and Sec24 are phosphorylated by the Hrr25 kinase, modifications that are proposed to help in the uncoating of COPII vesicles (Bhandari et al., 2013; Lord et al., 2011). This evidence can be used to propose that Erv14p is phosphorylated at the Golgi where it could help in controlling the directionality of packaging Erv14 into COPI or COPII vesicles for correct transport of its cargo proteins.

Our results provide evidence that leads us to suggest that Ser-134 acts in concert with the acidic amino acids (**E**SG**DD**) at the C-terminus for controlled transport of cargos. Due to the neutral pH of the ER lumen, the acidic amino acids may establish an electrostatic interaction with cargo proteins, allowing their loading into the COPII vesicles, where S134 would not play an important role. Arrival at the Golgi, as it has been described by Lord et al. (Lord et al., 2011), would lead to the phosphorylation of several COPII components for the uncoating of the vesicles and their fusion with the acceptor Golgi membrane, exposing the vesicle lumen to a more acidic environment that would release the cargo, expose the kinase domain of Erv14 to be phosphorylated, to prevent its interaction with the COPII subunits.

## Materials and Methods

### Yeast strains and growth media

The yeast strains are listed in Table S1. Yeast cell cultures were grown at 30°C in standard rich medium YPD: 1% yeast extract (Difco), 2% pectone (Difco) and 2% glucose (Sigma-Aldrich, Carlsbad, CA, USA) or selective medium YNB containing: 0.67% yeast nitrogen base without amino acids (Difco) and 2% glucose, supplemented with amino acids appropriated for auxotrophic growth. Amino acids were used at a concentration of 20 μg/ml (Sigma-Aldrich, Carlsbad, CA, USA). For the mbSUS assay, the synthetic complete medium was prepared as described (Lalonde et al., 2010) The *S. cerevisiae BYT45erv14Δ* mutant strain, which is highly sensitive to alkali metal cations by deletion of the cation exporters ENA1-5 and NHA1 was derived from BY4741 cells (Navarrete et al., 2010). The cargo receptor *erv14Δ* mutant and doubled mutants BY4741*pdr12Δerv14Δ* and BY4741*qdr2Δerv14Δ* were generated earlier (Rosas-Santiago et al., 2017). For testing sensitivity to sodium, 10-fold serial dilutions of cultures were applied to YNB plates supplemented with NaCl concentration from 0.4 M to 0.8 M; cell growth was recorded for 5 d. For complementation assays of the thermosensitive mutants *Sec23-1 erv14Δ, Sec16-2 erv14Δ*, and *Sec13-1 erv14Δ* serial dilutions of cultures were applied with a replica plater (Sigma-Aldrich, R2383) to YNB plates and growth at 23°C, 28°C or 30°C; cell growth was recorded for 3 d. To monitor growth rate, yeast cells were preinoculated in selective medium and the next day an OD_600_ of 0.01 was inoculated in YNB medium and growth was monitored every 2 h in spectrophotometer (Cary 60 UV-Vis, Agilent). Yeast cells were transformed using a standard lithium acetate transformation protocol (Gietz and Woods, 2006).

### Plasmid construction

To generate the point mutations S134A and S134D, the ORF of *ScERV14* was mutated by PCR using Phusion High-Fidelity DNA Polymerase (Thermo Fisher Scientific, Waltham, MA, USA) and employing the specific primers described in Table S2. The PCR products for *ERV14^S134A^* and *ERV14^S134D^* were cloned into the pDONOR207 vector of the Gateway system following manufacturer’s instructions (Invitrogen, Carlsbad, CA, USA). The entry clones thus generated were confirmed by restriction analysis with EcoRI and BgllI (Thermo Scientific Fisher, Waltham, MA, USA) and by sequencing. The LR clonase reaction (Invitrogen, Carlsbad, CA, USA) was used to transfer the genes *ERV14, ERV14^S134A^ and ERV14^S134D^* into the multicopy plasmid pAG425-3HA (plasmid #14250 Addgene), to generate the HA-tagged fusions pERV14-3HA, pERV14^S134A^-3HA and pERV14^S134D^-3HA. To complement *ERV14* function in the strains *BYT45erv14Δ*, BY4741*pdr12Δerv14Δ* and BY4741*qdr2Δerv14Δ* mutant strains, the genes *ERV14, ERV14^S134A^* and *ERV14^S134D^* were cloned into the pDR-F1_GW by an LR clonase reaction (Invitrogen, Carlsbad, CA, USA) generating the constructs pDR-F1-ERV14, pDR-F1-ERV14^S134A^ and pDR-F1-ERV14^S134D^. For the mbSUS assay constructs, the genes *ERV14, ERV14^S134A^* and *ERV14^S134D^* were transferred into the pXN32_GW (Nub clones) vector by an LR clonase reaction (Invitrogene, Carlsbad, CA, USA), generating the pNub-ERV14, pNub-ERV14^*S134A*^ and pNub-ERV14^*S134D*^. The pQDR2-Cub (Cub clone) construction used in this work was obtained from a previous work (Rosas-Santiago et al., 2017). To generated GFP-tagged mutant Erv14 proteins, the *ERV14* ORF was amplifying by PCR employing the primers described in Table 2, and the PCR products were cloned into the pGRU1 by homologous recombination in *S. cerevisiae* BW31a, leaving the *ERV14* ORF under the NHA1 promoter and in-frame with GFP, to yield the constructions pERV14^*S134A*^-GFP and pERV14^*S134D*^-GFP. The constructions were confirmed by restriction analysis with PstI (Thermo Scientific, Waltham, MA, USA) and by sequencing, using the oligos described in the Table S2. The pGRU1-originating plasmid pNHA-985-GFP, pDR12-GFP, pQDR2-GFP and pERV14-GFP used in this work were described earlier (Rosas-Santiago et al., 2017).

### Mating Based Split Ubiquiting System (mbSUS)

For test the binding of Erv14 mutants to their cargo proteins, the mbSUS assay was used (Lalonde et al., 2017;). The THY.AP4 (*MATa ura3, leu2*, lexA::LacZ::*trp1 lexA::HIS3 lexA::ADE2*) and THY.AP5 (*MATα URA3, leu2, trp1, his3 loxP::ade2*) yeast strains were transformed with the pQdr2-Cub (Cub clones) and pNub-Erv14^*S134A*^ or pNub-Erv14^*S134D*^ (Nub clones) constructions, respectively. Cells were allowed to mate on YPD medium for 24 h at 30°C, and then, diploid cells were selected on synthetic complete (SC) medium. For testing interactions, serial dilutions of cells were applied on SC medium supplemented with methionine concentration from 5 μM to 500 μM, cell growth was recorded for 3 d. The β-galactosidase activity of the lacZ (Fig. 3B), was revealed with 5-bromo-4-chloro-3-indolyl β-D galactoside (X-gal) (Sigma-Aldrich, Carlsbad, CA, USA). Diploid cells were grown on SC medium supplemented with 20 mg/L adenine hemisulfate, 20 mg/L histidine-HCl and then covered with an X-gal-agarose layer (0.5% agarose, 0.5 M phosphate buffer, pH 7.0, 0.1% SDS, 0.6 mg/ml X-gal). Blue staining was developed for 1-24 h.

### Live-Cell Imaging

To analyze the subcellular localization of the GFP-tagged proteins, cells were grown in liquid YNB medium to mid-log phase. A portion (1 ml) of the cultures was harvested, centrifuged at 4000 rpm for 5 min using an GS15R table-top centrifuge (Eppendorf) and cell pellets resuspended in (25 μl) of YNB medium. Five μl of resuspended cells were placed on a slide and were observed under an inverted multiphotonic confocal microscope (Olympus FV1000) equipped with a 60× 1.3 N.A. oil immersion objective. GFP fluorescence was visualized by excitation with a Multi-line Argon laser at 488 nm and spectral detector was set between 515/30 nm for emission. Images were processed with Image J (Schindelin et al., 2012).

### Preparation of Semi-intact cells and membrane fractionation

To maintain selective pressure on the plasmids, yeast cells were grown overnight in selective medium YNB and diluted to an OD_600_ of 0.1 in 50 ml of YPD medium the following morning, and then grown to approximately an OD_600_ of 1.0 and harvested by centrifugation in an Sorvall RC 6 Plus centrifuge (3500 rpm, 5 min, SS-34 rotor) at room temperature. The cell pellet was resuspended in buffer (100 mM Tris, pH 9.4, 5 mM DTT), incubated for 10 min at room temperature and centrifuged as before. The cells were resuspended and converted to spheroplast by digestion with (+)-lyticase buffer (20 mM Tris, pH 7.5, 0.7 M sorbitol, 0.5% glucose, 2 mM DTT and 1 mg/ml litycase. The cells were incubated at room temperature until the OD_600_ of a 1:100 dilution in H_2_O was less than 80% of the initial value and then spheroplasted cells were diluted into ice-cold (-)-lyticase buffer (20 mM Tris, pH 7.5, 0.7 M sorbitol, 0.5% glucose, 2 mM DTT)followed by centrifugation at 5000 rpm (Sorvall RC 6 Plus centrifuge with an SS-34 rotor) for 5 min at 4°C. Spheroplasts were resuspend in B88 buffer (20 mM HEPES, pH 6.8, 250 mM sorbitol, 150 mM KOAc, 5 mM MgOAc), aliquoted, frozen in liquid N_2_, and stored at −80°C. Spheroplasts were used as a membrane source in the vesicle budding assay. Isolation of ER and Golgi enriched fractions followed a similar procedure. After converting to spheroplasts, pelleted cells were resuspended in 2.5 ml JR lysis buffer (20 mM HEPES pH 7.4, 0.1 M sorbitol, 50 mM KOAc, 2 mM EDTA, 1 mM DTT, 1 mM PMSF) and lysed with six strokes of a Dounce homogenizer and centrifuged in an SS-34 rotor at 2000 rpm at 4°C for 5 min, to pellet nuclei and unbroken cells. The supernatant was recovered and centrifuged at 14,000 rpm for 10 min at 4°C in an Eppendorf 5417R centrifuge to generate the ER-enriched p13 pellet fraction. The supernatant was recovered and centrifuged at 60,000 rpm for 15 min at 4°C in an Optima TL micro-ultracentrifuges (TLA 100.3 rotor Beckman Coulter, Fullerton, CA) to obtain the Golgi-enriched p100 pellet fraction. The p13 and p100 pellets were resuspended in 100 μl JR lysis buffer and diluted 1:1 with 5X sample buffer (0.125 M Tris/HCl pH 6.8, 25% glycerol, 4% SDS and trace of bromophenol blue), boiled, and separated on 12% SDS–PAGE gels. Immunoblotting was conducted with polyclonal antibodies against Yet3 (ER marker) and Och1 (Golgi marker), that were used as fractionation controls; anti-GFP (Sigma-Aldrich, G1544, Lot 025M4754V) was used to monitor Erv14-GFP or Nha1-GFP.

### In vitro vesicle budding assay

COPII vesicle budding assay were performed as previously described (Barlowe et al., 1994). Washed semi-intact cell (SICs) were obtained from cells expressing either HA-tagged wild type (Erv14) or the mutant versions (Erv14^*S134A*^) or (Erv14S^*134D*^) in *BY4742Δerv14* cells and incubated in the presence or absence of purified COPII proteins (Sar1p, Sec23p/Sec24p and Sec13p/Sec31p), GTP and an 10X ATP regeneration system, an incubated at 25 °C for 30 min, for reconstitution of vesicle formation (Barlowe et al. 1994). A pool of 12.5 μl from ±COPII reactions was mixed with 200 μl of B88 buffer, which represented 10% of the total reaction (T) that served as a loading control. SIC’s were pelleted at 14,000 rpm for 3 min at 4 °C in an Eppendorf 5417R centrifuge. The supernatant containing vesicles was centrifuged at 60,000 rpm (Eppendorf 5417R centrifuge with a TLA100.3 rotor; Beckman Coulter, Fullerton, CA) to collect the membranes. Membrane pellets were resuspended in 25 μl of sample buffer 5X, and 8 or 10 μl aliquots were loaded on 14% polyacrylamide gels and immunoblotted for Sec61p, Erv41p, Coy1p or anti-HA for Erv14-3HA (WT) or mutants Erv14-^*S134D*^-3HA, Erv14-^*S134A*^-3HA detection. To estimate packaging efficiency, the intensity of the bands was quantified using GeneTools image analysis software (Syngene), where (T) represents 10% of the reaction and was used to compare that from the vesicles obtained from the different Erv14 isoforms.

### Erv14 phosphoproteomic analysis by tandem mass spectrometry (LC–MS/MS)

For sample preparation, three independent microsomes replicates (10 ug per replicate) were run in an SDS-PAGE. Protein spot from the gel that corresponded to the bands recognized by the HA antibody were sent to the Proteomics Facility at the Institut de Recherches Cliniques de Montréal, Canada. The samples were analyzed using Mascot software (Matrix Science, London, UK). Mascot was set up to search the Uniprot_S_cerevisiae_txid_4932 database. Scaffold (version Scaffold_4.2.1, Proteome Software Inc., Portland, OR) was used to validate LC-<u>MS/MS</u>-based peptide and protein identifications. Scaffold parameters were set to a minimum of two peptides per protein and accepted if they could be stablished with minimum probabilities of 99% at the protein level by the Peptide Prophet algorithm.

### SDS-PAGE and protein immunoblotting

Protein for Western blot analysis was precipitated by dilution of the samples 50-fold in 1:1 (v/v) ethanol:acetone and incubated overnight at −30 °C as described (Parry et al., 1989). Samples were then centrifuged at 13,000×g for 20 min at 4 °C using an F2402 rotor in a GS15R table-top centrifuge (Eppendorf). Pellets were air dried, resuspended with SDS-PAGE sample buffer 4X (125 mM Tris, pH 6.8, 25% glycerol, 4% SDS, 5% β-mercaptoethanol, bromophenol blue), and heated at 95 °C for 5 min before loading onto 12% (w/v) linear acrylamide mini-gels. For vesicle budding assay samples, pellets were resuspended in sample buffer 4X, boiled at 95 °C for 5 min, and resolved onto 14% (w/v) SDS-polyacrylamide gels. After protein separation, gels were either fixed and stained with Coomassie Brilliant Blue R 250 or electrophoretically transferred onto nitrocellulose membranes (Millipore) by standard methods. Following transfer, proteins were stained with Ponceau S protein stain (0.1% w/v in 1% v/v acetic acid for 30 s) to check for equal loading/transfer of proteins. Membranes were then blocked with TBS (100 mM Tris, 150 mM NaCl) containing 0.02% (w/v) Na-azide and 5% (w/v) fat-free milk powder (Nestle, Mexico) for 2 h at room temperature. Blocked membranes were incubated overnight with the appropriate primary antibodies, anti-HA (Sigma-Aldrich, H6908, Lot # 098M4812V), anti-phospho-serine (Sigma-Aldrich, P3430, Lot # 114M470V) anti-GFP (Sigma-Aldrich, G1544, Lot # 025M4754V), anti-Erv41 and anti-Och1 (Otte et al. 2001), or anti-Sec61 (Stirling et al. 1992). Primary antibodies were used at 1:1000 dilutions except for anti-HA, which was used at 1:6000 dilutions. For secondary antibodies, horseradish peroxidase–linked anti-rabbit and anti-mouse antibodies (GE Healthcare) were used at 1:10,000 dilutions. Blots were developed with two different reagents, chemiluminescent Luminata™ Crescendo procedure (Millipore) or SuperSignal West Pico Chemiluminescent substrate (Thermo Fisher Scientific, Waltham, MA, USA). Images were captured using either Gel DOC™ XR+ SYSTEM (BIORAD) or G:Box Chemi XR5 (Syngene).

### Analysis of phosphoproteins by Zn(II)-Phos-Tag SDS-PAGE

Phos-Tag SDS-PAGE protein separation was performed according to manufacturer’s specifications (Wako Pure Chemical Industries). The Phos-tag gel was prepared with 10% acrylamide and 50 μM Phos-tag as described here. For the resolving gel, 2.67 mL of (30% acrylamide/bis, 2.5 mL of 1.5 M Tris/HCl pH 8.8, 100 μL of 5 mM Phos-tag (Wako Pure Chemical Industries), 100 μL of 10 mM ZnCl_2_, 100 μL of 10% SDS, 10 μL TEMED, 25 μL of 10% APS and 4.5 ml of distilled water. The stacking gel was prepared with 600 μL of 30% acrylamide/bis-acrylamide (Sigma-Aldrich, Carlsbad, CA, USA), 1 mL of 0.5 M Tris/HCl pH 6.8, 40 μL of 10% SDS, 4 μL TEMED, 20 μL of 10% ammonium persulfate and 2.34 mL of distilled water. The gel was run at 30 mA for 6 h at 4 °C. After electrophoresis, the resolving gel was washed tree times in 100 mL of transfer buffer (192 mM glycine, 25 mM Tris-base pH 8.0, and 10% methanol containing 10 mM EDTA) and then washed once for 20 min in transfer buffer without EDTA. Protein was blotted onto nitrocellulose membrane at 30 V (at constant voltage) for 8 h at 4°C. Membranes were blocked using 5% nonfat milk and probed with 1:10000 dilution of anti-HA (Sigma-Aldrich, H6908, Lot # 098M4812V).

### Electron microscopy

For transmission electron microscopy, samples were prepared as described previously (Wright, 2000). Yeast cells were grown in YNB selective medium at mid-log phase (OD_600_ 0.8). Cells were centrifuged at 1500 g for 5 min in an F2402 rotor in a GS15R table-top centrifuge (Beckman) and resuspend with water to wash the cells. The pellet was fixed in ice-cold fixing Cacodylate buffer (0.1 M sodium cacodylate, pH 6.8; 1 mM MgCl_2_; 1 mM CaCl_2_ and 2% glutaraldehyde) for 2 h at 4°C and centrifuged as before. Cells were resuspended in Cacodylate buffer and incubated overnight at 4°C. Cells were pelleted as before and washed three times in Cacodylate buffer; then, cells were resuspended in 2% KMnO4 for 1 h at room temperature. After secondary fixing, cells were resuspended in 1 ml of freshly prepared 2% uranyl acetate for 1 h at room temperature and subsequently washed in water. Dehydration of cells was followed by clearing with propylene oxide and infiltration with a combination of 100% propylene oxide and 100% Spurr low-viscosity resin (TED PELLA.INC, USA). Finally, samples were embedded in 100% resin and kept in a vacuum oven at 45°C for 12 h, then shifted to 68°C for 3 d. For TEM examination, thin sections of grey-silver color interference (60–70 nm) were cut and mounted onto 300-mesh copper grids. The mounted sections were stained with alkaline lead citrate, washed gently with distilled water, and allowed to dry for 1 h. Dried sections were observed using a CARL ZEISS Libra 120 (Carl Zeiss AG, Oberkochen, Germany) transmission electron microscope, at a voltage of 80 kV, coupled with a Gatan Ultrascan 1000 CCD (2000 × 2000 pixels) (MultiScan camera model 794 gatan, Pleasanton, California, USA) to record the images.

## Acknowledgements

Assistance from Maria Guadalupe Muñoz Garcia in the lab is recognized. Assistance from the Laboratorio Nacional de Microscopia Avanzada (UNAM, México) is acknowledged. We also thank Jaime Blais and Kristofer J. Kaiser for their assistance. D. L-G, and C. Y-D were supported by CONACYT-México fellowships.

## Footnotes

### Author contributions

Conceptualization: D. L-G, O.P.; Performed the experiments: D. L-G., C. Y-D.; Performed the electron microscopy: G. Z-P.; Data analysis: D. L-G., C. Y-D., O.P.; Writing - original draft: D. L-G., O.P.; Writing - review & editing: D. L-G., O.P., C.B., C. Y-D., G. Z-P.; Supervision: O.P.; Funding acquisition: O.P.

## References

Anderson, N. S., Mukherjee, I., Bentivoglio, C. M. and Barlowe, C. (2017). The golgin protein Coy1 functions in intra-Golgi retrograde transport and interacts with the COG complex and Golgi SNAREs. Mol. Biol. Cell 28, 2686–2700.

Appenzeller, C., Andersson, H., Kappeler, F. and Hauri, H. P. (1999). The lectin ERGIC-53 is a cargo transport receptor for glycoproteins. Nat. Cell Biol. 1, 330–334.

Barlowe, C., Or, L., Yeung, T., Salama, N. and Rexach, M. F. (1994). COPII: A Membrane Coat Formed by Set Proteins That Drive Vesicle Budding from the Endoplasmic Reticulum. Cell 77, 895–907.

Belden, W. J. and Barlowe, C. (2001). Role of Erv29p in Collecting Soluble Secretory Proteins into ER-Derived Transport Vesicles. Science (80-.). 294, 1528–1531.

Bhandari, D., Zhang, J., Menon, S., Lord, C., Chen, S., Helm, J. R., Thorsen, K., Corbett, K. D., Hay, J. C. and Ferro-Novick, S. (2013). Sit4p/PP6 regulates ER-to-Golgi traffic by controlling the dephosphorylation of COPII coat subunits. Mol. Biol. Cell 24, 2727–2738.

Brandizzi, F. and Barlowe, C. (2013). Organization of the ER-Golgi interface for membrane traffic control. Nat. Rev. Mol. Cell Biol. 14, 382–92.

Dancourt, J. and Barlowe, C. (2010). Protein sorting receptors in the early secretory pathway. Annu. Rev. Biochem. 79, 777–802.

Davis, S., Wang, J., Zhu, M., Stahmer, K., Lakshminarayan, R., Ghassemian, M., Jiang, Y., Miller, E. a. and Ferro-Novick, S. (2016). Sec24 phosphorylation regulates autophagosome abundance during nutrient deprivation. Elife 5, 1–22.

Gietz, R. D. and Woods, R. a. (2006). Yeast transformation by the LiAc/SS Carrier DNA/PEG method. Methods Mol. Biol. 313, 107–120.

Grefen, C., Obrdlik, P. and Harter, K. (2009). The determination of protein-protein interactions by the mating-based split-ubiquitin system (mbSUS). Methods Mol. Biol. 479, 217–233.

Herzig, Y., Sharpe, H. J., Elbaz, Y., Munro, S. and Schuldiner, M. (2012). A systematic approach to pair secretory cargo receptors with their cargo suggests a mechanism for cargo selection by Erv14. PLoS Biol. 10, e1001329.

Kaiser, C. A. and Schekman, R. (1990). Distinct sets of SEC genes govern transport vesicle formation and fusion early in the secretory pathway. Cell 61, 723–733.

Kinoshita, E. and Kinoshita-Kikuta, E. (2011). Improved Phos-tag SDS-PAGE under neutral pH conditions for advanced protein phosphorylation profiling. Proteomics 11, 319–323.

Koreishi, M., Yu, S., Oda, M., Honjo, Y. and Satoh, A. (2013). CK2 Phosphorylates Sec31 and Regulates ER-To-Golgi Trafficking. PLoS One 8, 1–9.

Lord, C., Bhandari, D., Menon, S., Ghassemian, M., Nycz, D., Hay, J., Ghosh, P. and Ferro-Novick, S. (2011). Sequential interactions with Sec23 control the direction of vesicle traffic. Nature 473, 181–186.

Martínez-Menárguez, J. a., Geuze, H. J., Slot, J. W. and Klumperman, J. (1999). Vesicular tubular clusters between the ER and Golgi mediate concentration of soluble secretory proteins by exclusion from COPI-coated vesicles. Cell 98, 81–90.

Meggio, F. and Pinna, L. A. (2003). One-thousand-and-one substrates of protein kinase CK2? FASEB 17, 349–368.

Miller, E., Antonny, B., Hamamoto, S. and Schekman, R. (2002). Cargo selection into COPII vesicles is driven by the Sec24p subunit. EMBO J. 21, 6105–6113.

Muñiz, M., Nuoffer, C., Hauri, H. P. and Riezman, H. (2000). The Emp24 complex recruits a specific cargo molecule into endoplasmic reticulum-derived vesicles. J. Cell Biol. 148, 925–930.

Nakagawa, T. (2019). Structures of the AMPA receptor in complex with its auxiliary subunit cornichon. Science (80-.). 366, 1259–1263.

Navarrete, C., Petrezsélyová, S., Barreto, L., Martínez, J. L., Zahrádka, J., Ariño, J., Sychrová, H. and Ramos, J. (2010). Lack of main K+ uptake systems in Saccharomyces cerevisiae cells affects yeast performance in both potassium-sufficient and potassium-limiting conditions. FEMS Yeast Res. 10, 508–517.

Otte, S., Belden, W. J., Heidtman, M., Liu, J., Jensen, O. N. and Barlowe, C. (2001). Erv41p and Erv46p: New components of COPII vesicles involved in transport between the ER and Golgi complex. J. Cell Biol. 153, 503–517.

Pagant, S., Wu, A., Edwards, S., Diehl, F. and Miller, E. a. (2015). Sec24 is a coincidence detector that simultaneously binds two signals to drive ER export. Curr. Biol. 25, 403–412.

Park, B. C., Reese, M. and Tagliabracci, V. S. (2019). Thinking outside of the cell: Secreted protein kinases in bacteria, parasites, and mammals. IUBMB Life 71, 749–759.

Parry, R. V., Turner, J. C. and Rea, P. a. (1989). High purity preparations of higher plant vacuolar H+-ATPase reveal additional subunits. Revised subunit composition. J. Biol. Chem. 264, 20025–20032.

Piper, P., Mahé, Y., Thompson, S., Pandjaitan, R., Holyoak, C., Egner, R., Mühlbauer, M., Coote, P. and Kuchler, K. (1998). The Pdr12 ABC transporter is required for the development of weak organic acid resistance in yeast. EMBO J. 17, 4257–4265.

Powers, J. and Barlowe, C. (1998). Transport of Axl2p Depends on Erv14p, an ER–Vesicle Protein Related to the. J. Cell Biol. 142, 1209–1222.

Powers, J. and Barlowe, C. (2002). Erv14p Directs a Transmembrane Secretory Protein into COPII-coated Transport Vesicles. Mol. Biol. Cell 13, 880–891.

Rosas-Santiago, P., Zimmermannova, O., Vera-Estrella, R., Sychrová, H. and Pantoja, O. (2016). Erv14 cargo receptor participates in yeast salt tolerance via its interaction with the plasma-membrane Nha1 cation/proton antiporter. Biochim. Biophys. Acta - Biomembr. 1858, 67–74.

Rosas-Santiago, P., Lagunas-Gomez, D., Yáñez-Domínguez, C., Vera-Estrella, R., Zimmermannová, O., Sychrová, H. and Pantoja, O. (2017). Plant and yeast cornichon possess a conserved acidic motif required for correct targeting of plasma membrane cargos. Biochim. Biophys. Acta - Mol. Cell Res. 1864, 1809–1818.

Salvi, M., Sarno, S., Cesaro, L., Nakamura, H. and Pinna, L. A. (2009). Extraordinary pleiotropy of protein kinase CK2 revealed by weblogo phosphoproteome analysis. Biochim. Biophys. Acta - Mol. Cell Res. 1793, 847–859.

Sato, K. and Nakano, A. (2007). Mechanisms of COPII vesicle formation and protein sorting. FEBS Lett. 581, 2076–2082.

Schindelin, J., Arganda-Carreras, I., Frise, E., Kaynig, V., Longair, M., Pietzsch, T., Preibisch, S., Rueden, C., Saalfeld, S., Schmid, B., et al. (2012). Fiji: An open-source platform for biological-image analysis. Nat. Methods 9, 676–682.

St-Denis, N., Gabriel, M., Turowec, J. P., Gloor, G. B., Li, S. S. C., Gingras, A. C. and Litchfield, D. W. (2015). Systematic investigation of hierarchical phosphorylation by protein kinase CK2. J. Proteomics 118, 49–62.

Stagljar, I., Korostensky, C., Johnsson, N. and Te Heesen, S. (1998). A genetic system based on split-ubiquitin for the analysis of interactions between membrane proteins in vivo. Proc. Natl. Acad. Sci. U. S. A. 95, 5187–5192.

Thor, F., Gautschi, M., Geiger, R. and Helenius, A. (2009). Bulk flow revisited: Transport of a soluble protein in the secretory pathway. Traffic 10, 1819–1830.

Vargas, R. C., Tenreiro, S., Teixeira, M. C., Fernandes, A. R. and Sá-Correia, I. (2004). Saccharomyces cerevisiae multidrug transporter Qdr2p (Yil121wp): Localization and function as a quinidine resistance determinant. Antimicrob. Agents Chemother. 48, 2531–2537.

Wright, R. (2000). Transmission electron microscopy of yeast. Microsc. Res. Tech. 51, 496–510.

Zanetti, G., Pahuja, K. B., Studer, S., Shim, S. and Schekman, R. (2012). COPII and the regulation of protein sorting in mammals. Nat. Cell Biol. 14, 221.

