## Supplemental Information for "The C-terminus of the cargo receptor Erv14p affects COPII vesicle formation and cargo delivery"

**A**

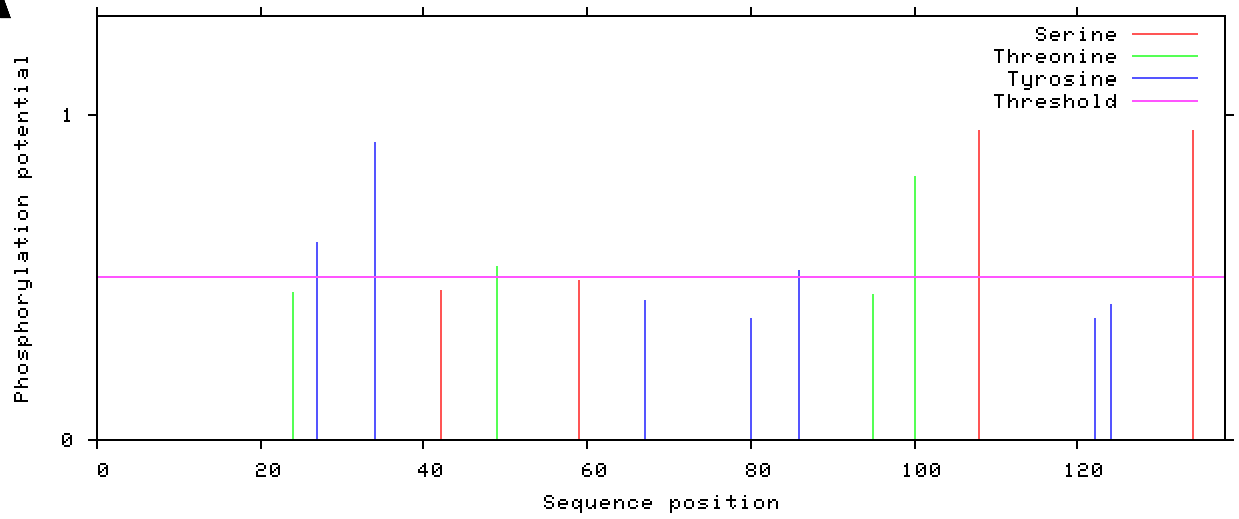


**B**

| #Aminoacid | Context | Score |
|---|---|---|
| **108S** | **HKRESFLKL** | **0.949** |
| **134S** | **LIAESGDDF** | **0.948** |

**Supplementary Figure 1. Search of putative phosphorylation sites using Netphos 3.1. A).** The plot shows predicted phosphorylation sites in the Erv14 sequence. **B).** The two possible phosphorylatable Serine residues and their sequence consensus.

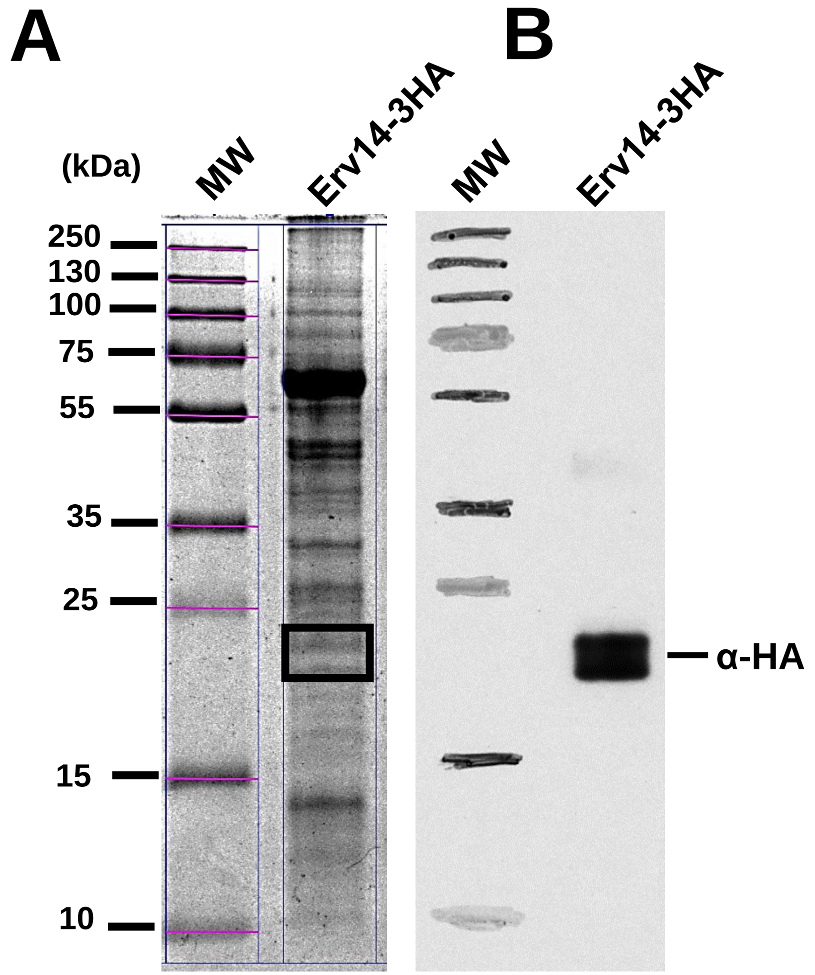


**C**

| Sample | Protein Name | Protein MW (kDa) | Unique Peptide Count | Unique Spectra Count | Total Ion Spectra |
|---|---|---|---|---|---|
| **1** | **Erv14** | 16 | 2 | 3 | 9 |
| **2** | **Erv14** | 16 | 2 | 2 | 6 |
| **3** | **Erv14** | 16 | 2 | 2 | 6 |

**D**

| Calc. Mass (Da) | Obser. Mass (m/z) | ± Da | Start Seq. | End Seq. | Sequence | Ion Score |
|---|---|---|---|---|---|---|
| **1822.00** | 68.34 | 0.0036 | 85 | 99 | (K)IYNKVQLLDATEIFR(T) | 30.7 |
| **1303.71** | 682.86 | 0.0011 | 89 | 99 | (K)VQLLDATEIFR(T) | 31.6 |

**Supplementary Figure 2. Search for phosphorylation sites by LC-MS/MS. A)** SDS-PAGE gel showing the selected area (square). **B)** Western blot developed with (α-HA) antibody. **C)** Table showing the number of peptides identified and the number of total spectra obtained by LC-MS/MS. **D)** Properties of the identified peptides.

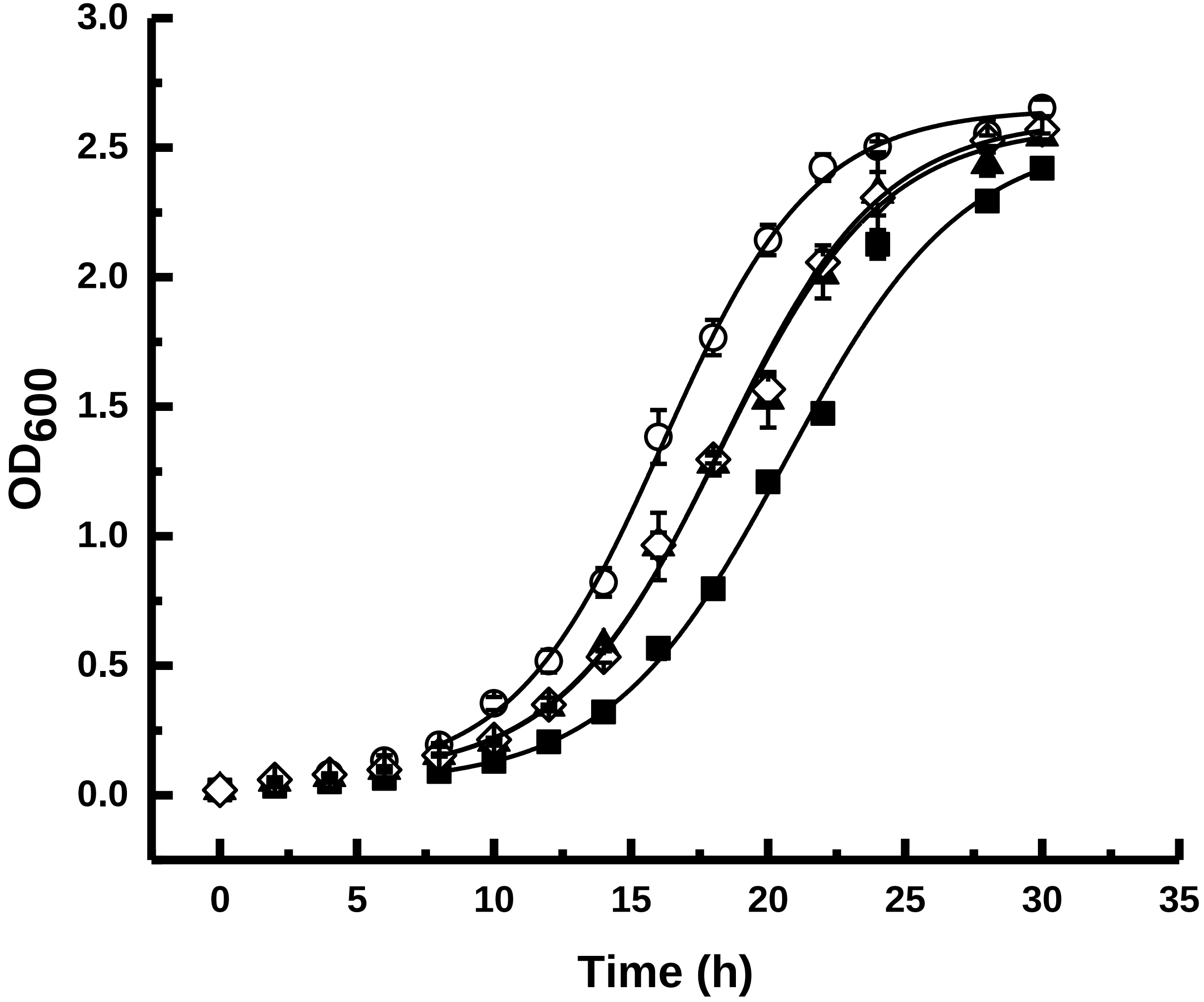


**Supplementary Figure 3. Cell growth curve from BY4741*erv14Δ* strain expressing wild type Erv14 and mutants.**

(○) Wild type Erv14, (▲) Dephosphorylated form Erv14$^{S134A}$, (◊) Phosphomimetic form Erv14$^{S134D}$, (■) Empty vector pDR-F1.

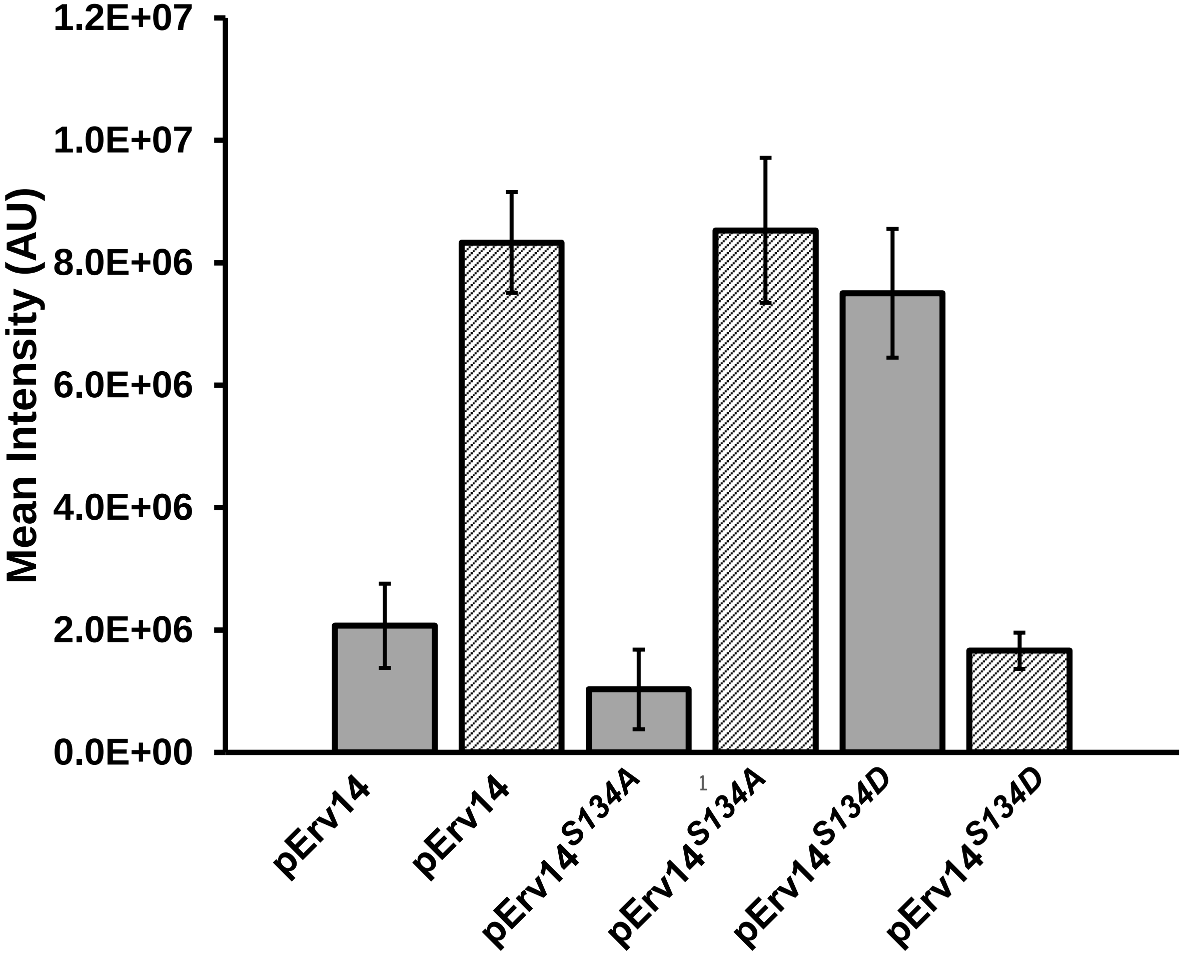


**Supplementary Figure 4. Quantification of band intensities from Figure 1B.** Intensities from the bands corresponding to the recognition of Nha1-GFP in the ER (grey bars) or PM/GA (hatched bars) by the anti-GFP antibody from yeast cells co-transformed with the *ERV14* or the corresponding mutant. Data are the mean ± SD from three different preparations.

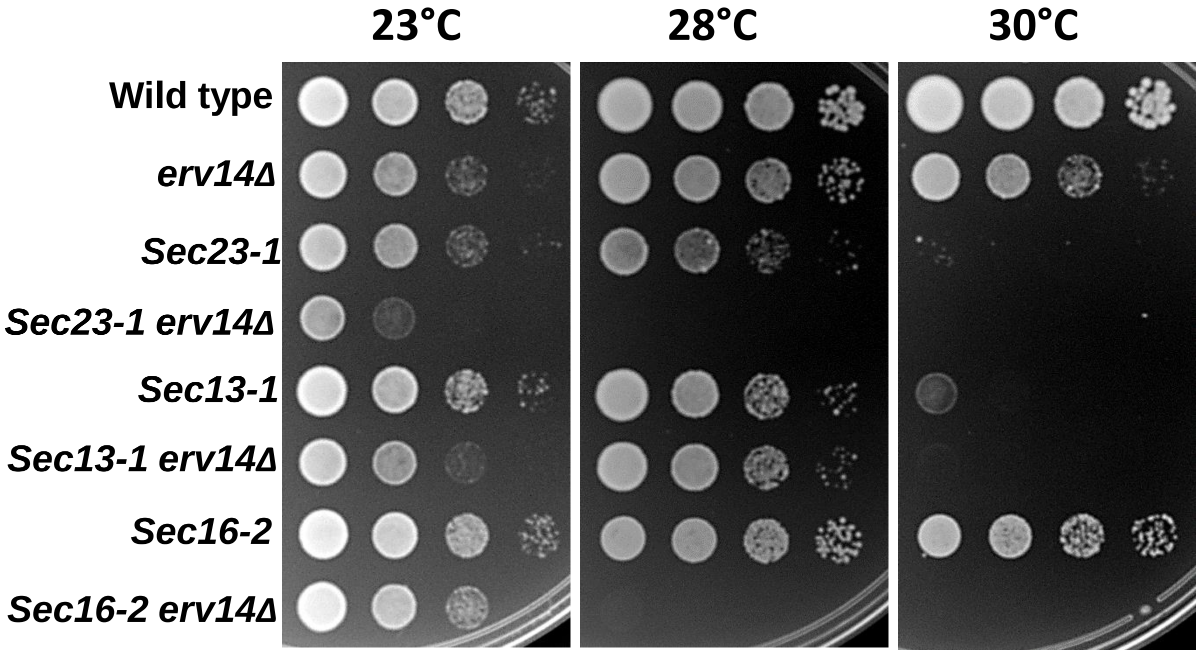


**Supplementary Figure 5. Genetic analysis of *erv14Δ* on *sec* mutants**. Disruption of *ERV14* increased the thermosensitivity of the *sec23-1, sec13-1* or *sec16-2* strains. Serial dilutions of the thermosensitivity strains were tested for growth at 23-30°C.

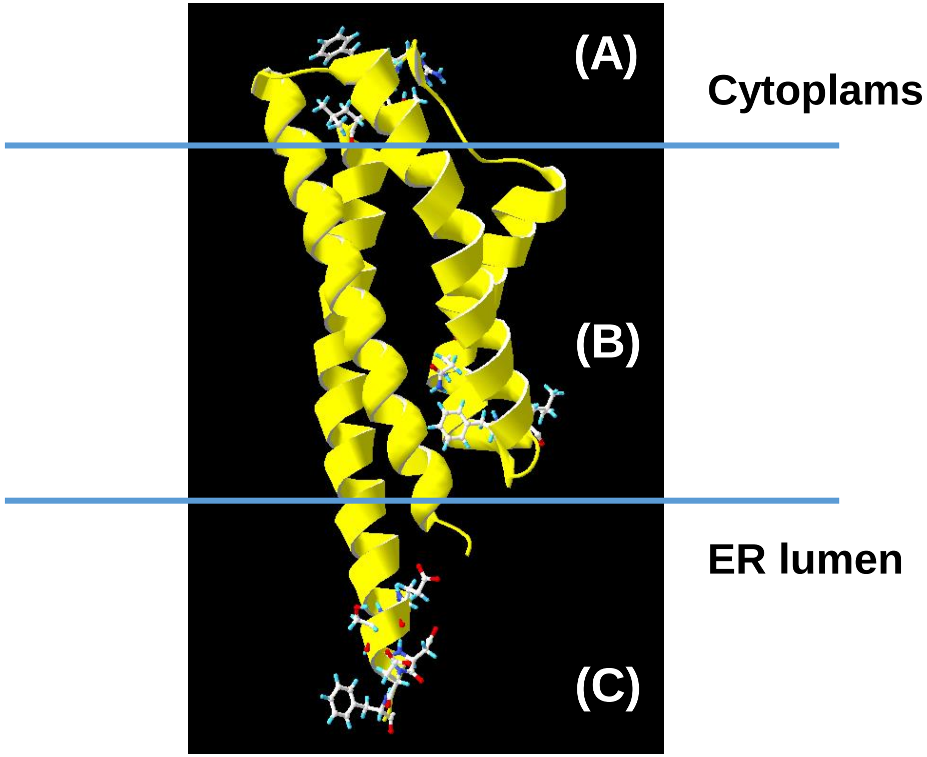


**Supplementary Figure 6. Erv14/cornichon molecular model obtained with the Alphafold2 algorithm.** Erv14 was predicted to possess four membrane spanning helices with the C and N termini facing the ER lumen. **A**) The COPII-binding site (IFRTL); **B**) Cargo-binding site; **C**) Possible phosphorylation domain. The position of the membrane limits is approximate

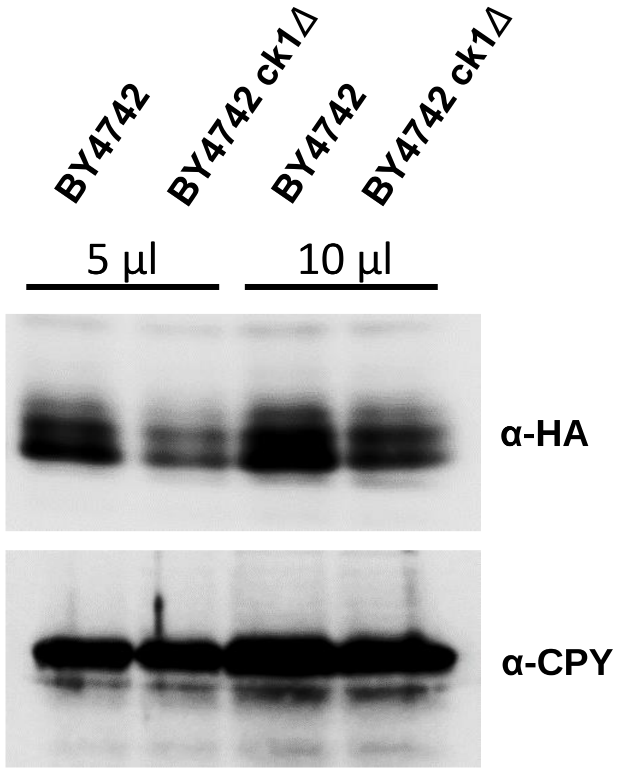


**Supplementary Figure 7. Analysis of Erv14 isoforms on the *ck1Δ* mutant.** Western blot demonstrating the abundance of Erv14-3HA in the microsomal fraction from BY4742 or BY4742*ck1Δ* cells. Observe the lower abundance of Erv14 when expressed in BY4742*ck1Δ.* CPY was used as a loading control.

**Fig. S8. Blot Transparency**

#### Related to Figure 1

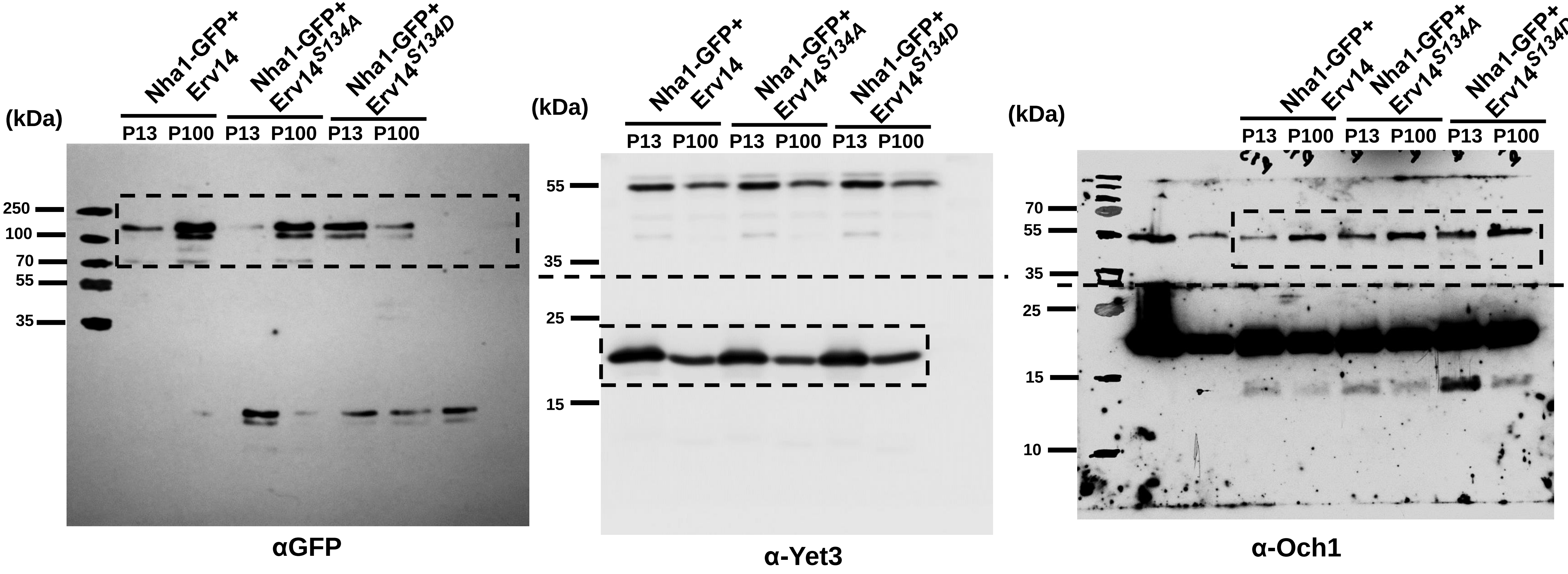


**Blots from plasma membrane localization of the Nha1 exchanger in BYT45*erv14Δ* cells transformed with Wild type pERV14 and point mutations**. Immunoblotting was conducted with Anti-GFP to monitor Nha-GFP (130 kDa). Yet3 (P13) and Och1 (P100) were used as fractionation controls. Dashed line and inset indicated cut regions shown in Figure 1.

#### Related to Figure 4

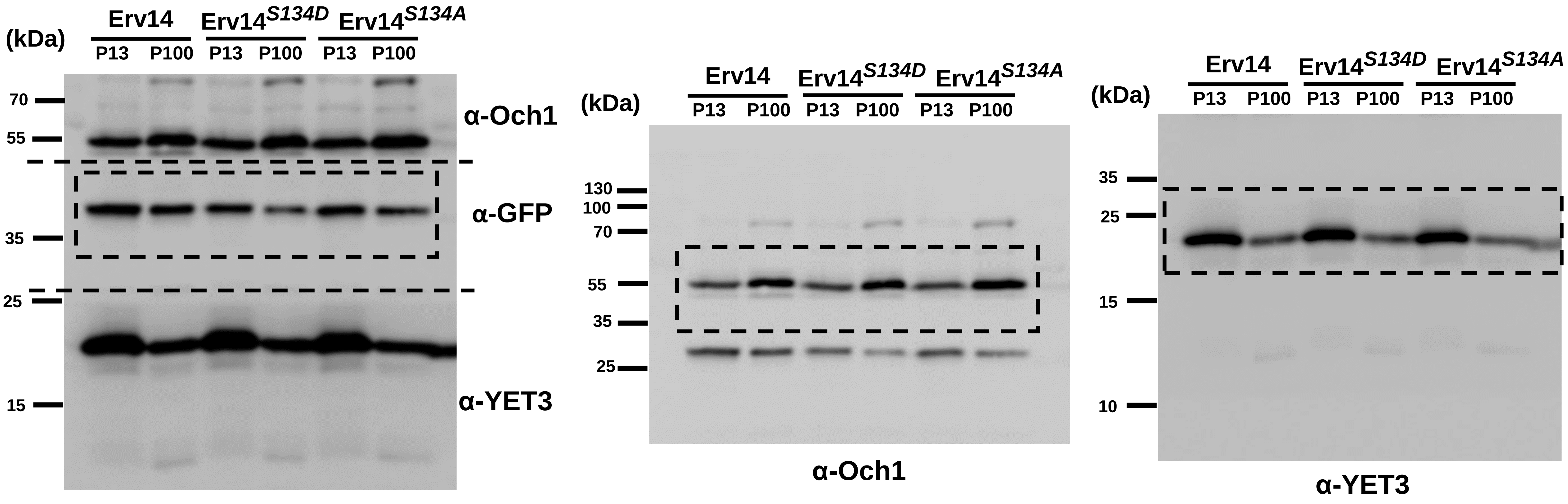


**Blots from Erv14 membrane distribution in ER or Golgi fractions from GFP-tagged wild-type and Erv14$^{S134A}$ or Erv14$^{S134D}$ mutants.** Immunoblotting was conducted with polyclonal antibodies against Yet3 (P13) and Och1 (P100), that were used as fractionation controls; anti-GFP was used to monitor Erv14-GFP (44 kDa). Dashed line and inset indicated cut regions shown in Figure 4.

#### Related to Figure 6

Blots from wild-type cargo receptor (Erv14-3HA), or the mutants (Erv14$^{S134A}$-3HA) or (pErv14$^{S134D}$-3HA) were incubated with (+) or without (-) purified COPII proteins to monitor vesicle budding. (T) represent total budding reaction. Anti-HA was used to monitor Erv14-3HA (19 kDa), Erv41 (40 kDa) or Coy1(77 kDa) antibodies was used as a positive control, Sec61 (53 kDa) as a negative control. Dashed line and inset indicate cut regions shown in Figure 6.

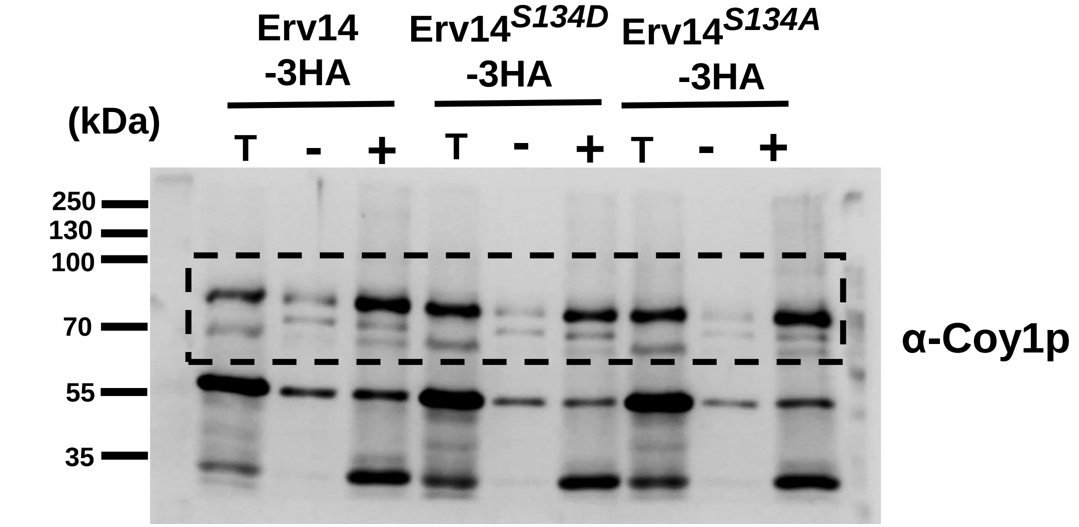


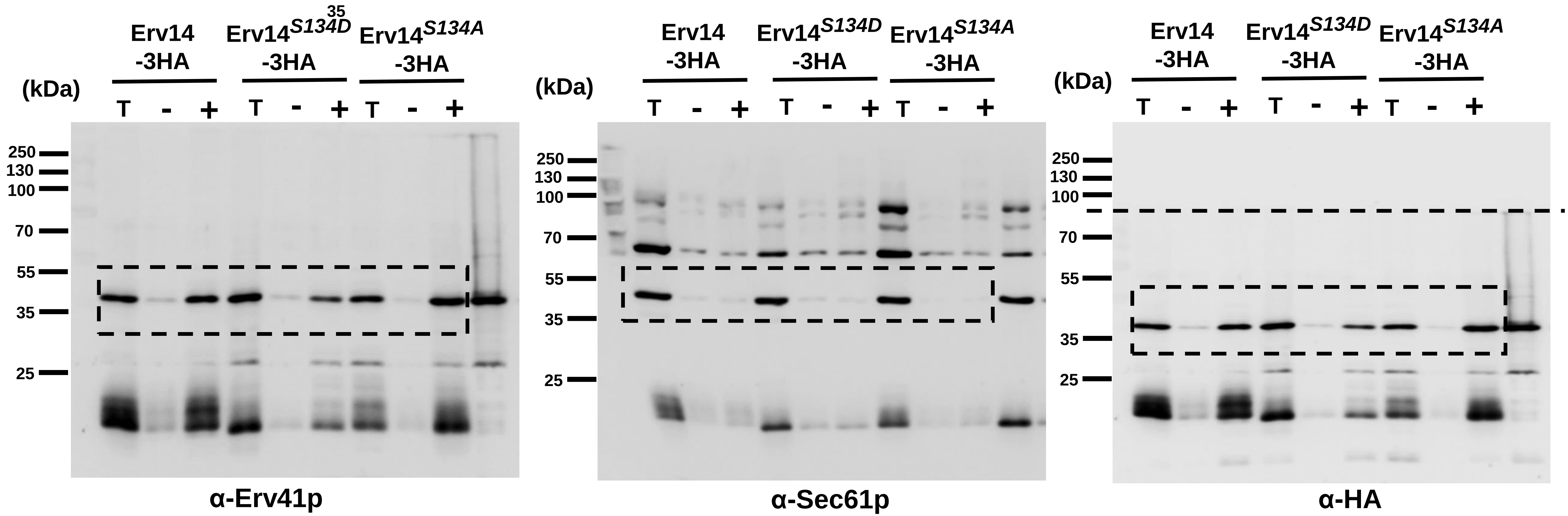

### Related to Figure 7

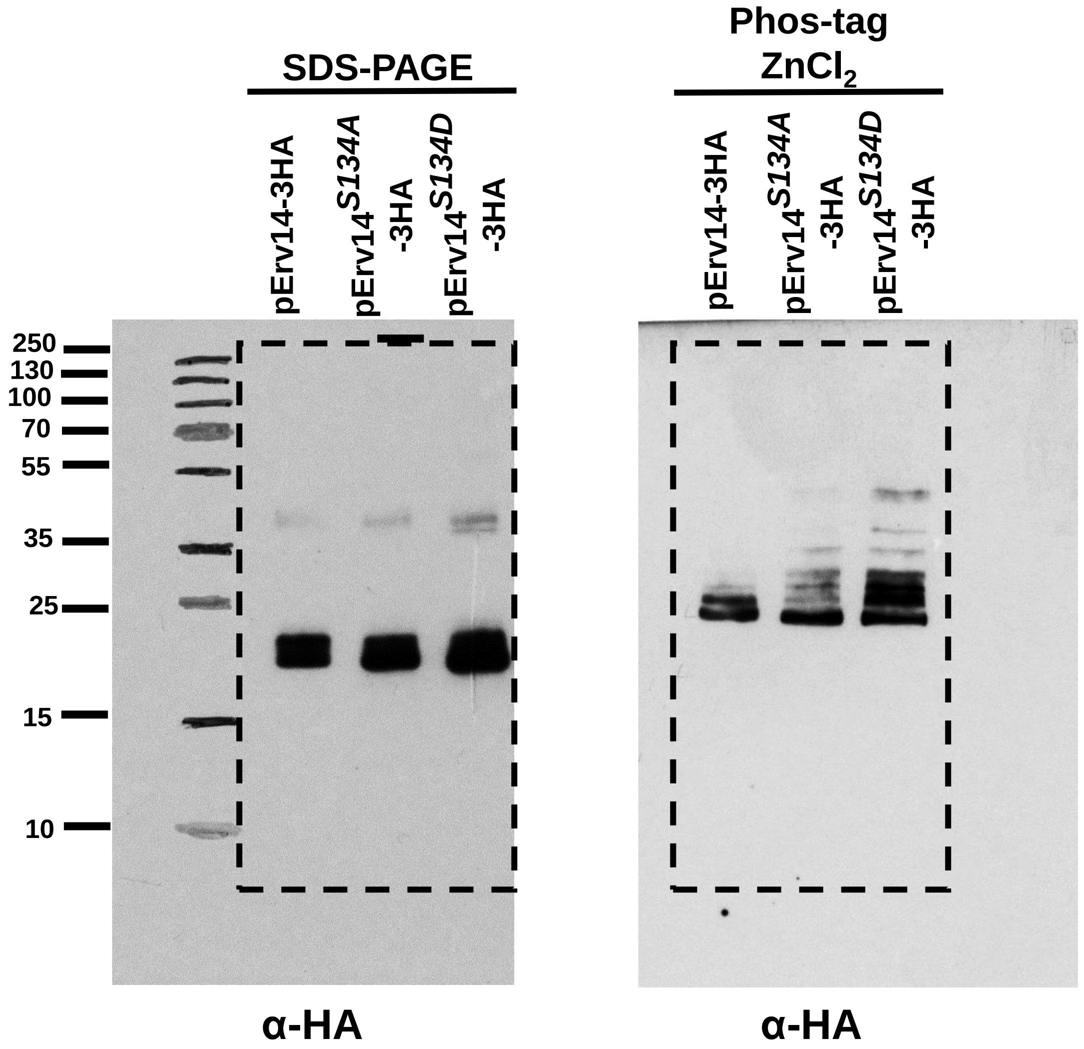


Blots from HA-tagged Erv14, Erv14$^{S134A}$ or Erv14$^{S134D}$ expressing cells were separated in SDS-PAGE (left) or Zn(II)-Phos-tag (right) gels. Anti-HA antibody (19 kDa) was used to developed. Dashed inset indicated the cut region used in Figure 7

### Table S1 List of strains used in this study

| Strain | Genotype | Source |
|---|---|---|
| BY4741Δerv14 | MATa his3Δ1 leu2Δ0 met15Δ0 ura3Δ0 erv14Δ::KanMx | |
| BY4741Δerv14Δpdr12 | MATa his3Δ1 leu2Δ0 met15Δ0 ura3Δ0 erv14Δ::loxP pdr12Δ::KanMx | |
| BY4741Δerv14Δqdr2 | MATa his3Δ1 leu2Δ0 met15Δ0 ura3Δ0 erv14Δ::loxP qdr2Δ::KanMx | |
| *BYT45erv14Δ* | MATa his3Δleu2Δmet15Δura3Δ; nha1Δ::loxP ena1-5Δ::loxP erv14Δ::KanMx | |
| BY4742Δerv14 | MATα his3Δ1 leu2Δ0 lys2Δ0 ura3Δ0 erv14Δ::KanMx | Barlowe Lab |
| CBY617 (FY834) | MATα his3Δ200 ura3-52 lys2Δ202 trp1Δ63 erv14::HIS3 sec23-1 | Barlowe Lab |
| CBY663 (FY834) | MATα his3Δ200 ura3-52 leu2Δ1 lys2Δ202 erv14::HIS3 sec16-2 | Barlowe Lab |
| CBY603 (FY834) | MATα his3Δ200 ura3-52 leu2Δ1 lys2Δ202 trp1Δ63 erv14::HIS3 sec13-1 | Barlowe Lab |
| THY.AP4 | MATa ura3 leu2 lexA::lacZ::trp1 lexA::HIS3 lexA::ADE2 | Frommer Lab |
| THY.AP5 | MATα URA3 leu2 trp1 his3 loxP::ade2 | Frommer Lab |
| BW31a | *MATa leu2-3/122 ura3-1 trp1-1 his3-11/15 ade2-1 can1-100 GAL SUC2 mal10 ena1- 4Δ::HIS3 nha1::LEU2* | |

**Table S2**. Primers used for gene cloning into the Gateway vectors and pGRU1

| | 5´ | 3´ |
|---|---|---|
| **Gene** | **Forward** | **Reverse** |
| **ERV14$^{S134A}$-pDONOR221** | GTACAAAAAAGCAGGCTTCATGGGTGCTTGGTTATTTATC | GTACAAGAAAGCTGGGTCGAAGTCATCACCAGCTTCAGCAATC |
| **ERV14$^{S134D}$-pDONOR207** | GTACAAAAAAGCAGGCTTCATGGGTGCTTGGTTATTTATC | GTACAAGAAAGCTGGGTCGAAGTCATCACCATCTTCAGCAATC |
| **ERV14$^{S134A}$-pGRU1** | GTACATTATAAAAAAAAATCCTGAACTTAGCTAGATATTATGGGTGCTTGGTTATTTATCC | TAAAGCTCCGGAGCTTGCATGCCTGCAGGTCGACTCTGAAGTCATCACCAGCTTCAGCAATC |
| **ERV14$^{S134D}$-pGRU1** | GTACATTATAAAAAAAAATCCTGAACTTAGCTAGATATTATGGGTGCTTGGTTATTTATCC | TAAAGCTCCGGAGCTTGCATGCCTGCAGGTCGACTCTGAAGTCATCACCATCTTCAGCAATC |
